# Glutamate-Handling Proteins Associate with Adverse Clinicopathologic Features and Comorbidities in Invasive Lobular Carcinoma

**DOI:** 10.1101/2024.09.29.615681

**Authors:** Shaymaa Bahnassy, Todd A. Young, Theresa C. Abalum, Eden A. Pope, Amanda Torres Rivera, Aileen I. Fernandez, Ayodeji O. Olukoya, Dua Mobin, Suman Ranjit, Nicole E. Libbey, Sonali Persaud, Aaron M. Rozeboom, Krysta Chaldekas, Brent T. Harris, Zeynep Madak-Erdogan, Joseph L. Sottnik, Matthew J. Sikora, Rebecca B. Riggins

## Abstract

Invasive Lobular Carcinoma (ILC) is a subtype of breast cancer characterized by distinct biological features, and limited glucose uptake coupled with increased reliance on amino acid and lipid metabolism. Our prior studies highlight the importance of glutamate as a key regulator of ILC tumor growth and therapeutic response. Here we examine the expression of four key proteins involved in glutamate transport and metabolism – SLC3A2, SLC7A11, GPX4, and GLUD1/2 – in a racially diverse cohort of 72 estrogen receptor-positive (ER+) ILC and 50 ER+ invasive ductal carcinoma, no special type (IDC/NST) patients with primary disease. All four proteins associate with increased tumor size in ILC, with three showing stronger associations in Black women, but not in IDC/NST. Among these three proteins in ILC, GLUD1/2 uniquely associates with ER expression in all women, while GLUD1/2 and SLC3A2 are enriched in hypertensive women. GLUD1/2 and GPX4 are upregulated in endocrine therapy-resistant ILC cell lines, and pharmacological inhibition of GLUD1 reduces ER protein levels and cell viability. Together, these findings support a potentially important role for glutamate metabolism in ILC and suggest GLUD1 and other glutamate-handling proteins as candidate targets for therapeutic intervention in ILC.

## Introduction

ILC (Invasive Lobular Carcinoma) is a special histological subtype of breast cancer that accounts for 10-15% of cases diagnosed annually^1,2^. ILC has unique genetic, transcriptomic, and biological features as compared to the more common invasive ductal breast cancer, or breast cancer of no special type (IDC/NST). Nearly all ILC is estrogen receptor-positive (ER+), and characterized by mutational inactivation of *CDH1*, the gene encoding E-cadherin^3^, which is associated with the diffuse growth pattern of ILC tumors. Additionally, ILC metastasizes to the lung, bone, and brain like other ER+ breast malignancies, but it also has a propensity to spread to the digestive tract, ovaries, and the peritoneal cavity^1,2^. Complicating this atypical dissemination pattern, advanced or metastatic ILC may not be as easily detected by positron emission tomography using the glucose analog tracer ^18^F-fluorodeoxyglucose (i.e. [^18^F]FDG-PET)^4^, as several clinical studies have shown ILC has limited FDG avidity. While these observations complicate care of patients with metastatic ILC, they also suggest that ILC has a distinct metabolic phenotype among breast cancers that is less reliant on glucose uptake and metabolism.

Other PET tracers provide opportunities to understand differential ILC metabolism. Fluciclovine is an amino acid analog of leucine^5^, and prospective clinical trials demonstrate that ILC shows substantially higher [^18^F] Fluciclovine-PET uptake compared to [^18^F]FDG-PET]^6–8^. Among ILC, 100% of lesions were fluciclovine-positive while only 43% of lesions were FDG-positive. These findings are consistent with clinical and preclinical research, in which we and others show that ILC is more dependent on lipid and amino acid metabolism compared to glucose metabolism^9–12^. Our work suggests that the amino acid glutamate may be particularly important in ILC. Endocrine therapy-resistant ILC models upregulate multiple metabotropic glutamate receptors (mGluRs)^13^, and the small molecule Riluzole (a modulator of glutamate metabolism FDA approved for amyotrophic lateral sclerosis) is efficacious against *in vitro, in vivo*, and *ex vivo* models of ILC^14,15^, (discussed in^16^). Therefore, understanding glutamate metabolism in ILC can provide important opportunities to advance imaging and clinical management, as well as precision treatment paradigms.

For this study, we assembled a cohort of 72 ER+ primary ILCs and 50 ER+ primary IDC/NSTs with well-annotated clinical, pathological, and demographic data, and used multiplex immunohistochemistry (IHC) to determine the prognostic value of four key proteins in glutamate transport and metabolism (**Figure 1A**). Solute carriers (SLC) 3A2 (SLC3A2 or CD98) and 7A11 (SLC7A11) heterodimerize to form System xCT, an antiporter that imports cystine and exports glutamate. Downstream of xCT, glutathione peroxidase 4 (GPX4) protects cells from lipid peroxidation by catalyzing the reduction of lipid hydroperoxides and converting the glutamate-glycine-cysteine tripeptide glutathione (GSH) to glutathione disulfide (GSSG), thereby inhibiting ferroptosis (an iron-dependent mechanism of cell death). Finally, glutamate dehydrogenase 1/2 (GLUD1/2 or GDH1/2) is a bidirectional metabolic enzyme that in the forward direction converts glutamate to alpha-ketoglutarate to replenish the tricarboxylic acid (TCA) cycle, while in the reverse direction it recycles ammonia to bolster tumor cell growth and amino acid synthesis^17,18^.

**Figure 1.**
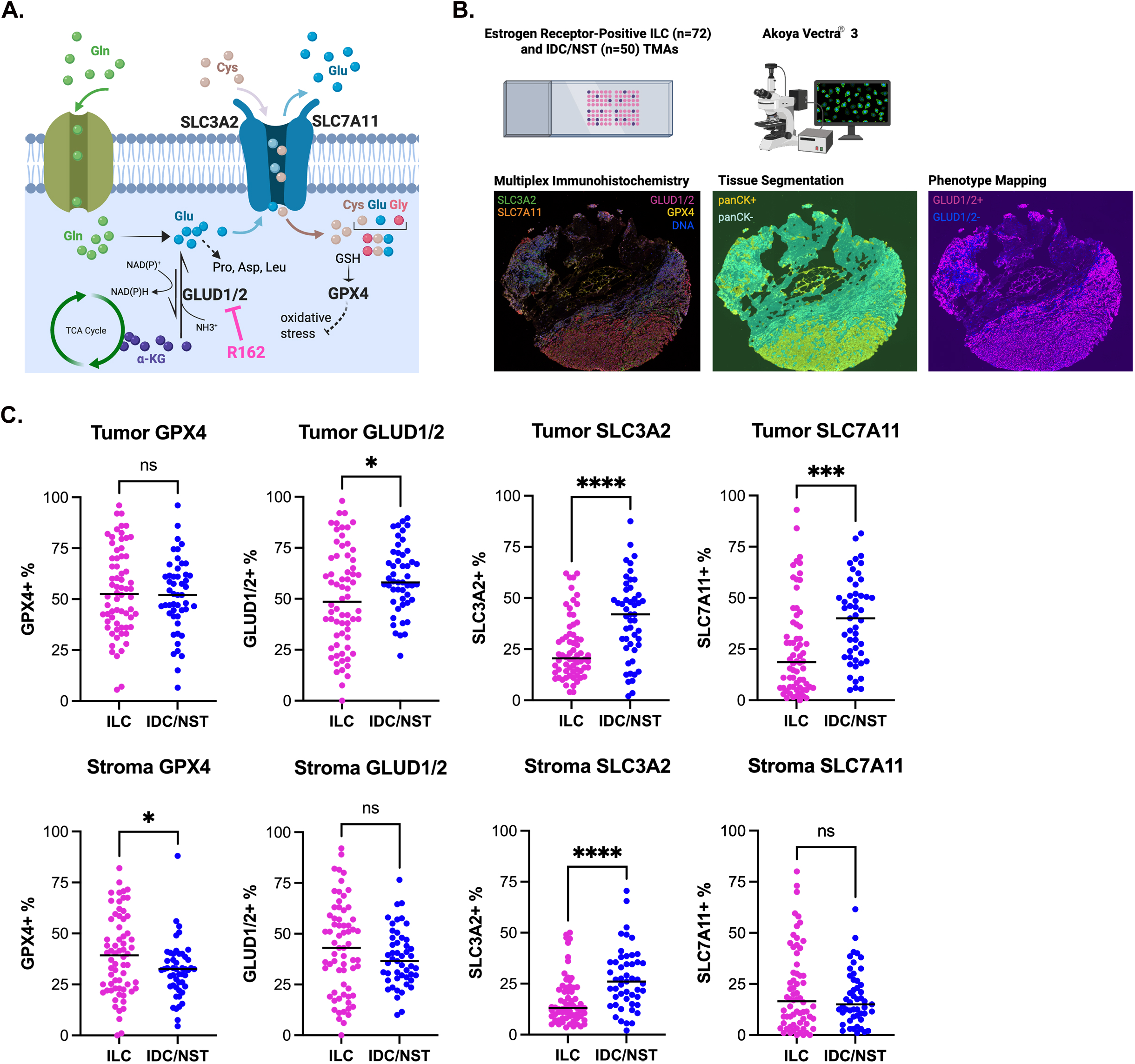
Expression of glutamate-handling proteins in ILC and IDC/NST. **A**, Schematic showing functional relationships between the four glutamate-handling proteins included in our multiplex IHC panel. Gln, glutamine; Glu, glutamate; Cys, cyst(e)ine; Gly, glycine; Pro, proline; Asp, aspartic acid; Leu, leucine; GSH, glutathione; R162, GLUD1 pharmacological inhibitor. **B**, Schematic showing cohort assembly and study workflow. A representative TMA core stained for the four target proteins and nuclear staining (DNA) is shown, along with its corresponding images processed for tissue segmentation, indicating areas of panCK+ (yellow, epithelial) and panCK-(aqua, stromal) cells, followed by phenotype mapping for (as an example) GLUD1/2+ (magenta) and GLUD1/2- (blue) cells. **C**, Comparison of protein expression for GPX4, GLUD1/2, SLC3A2 and SLC7A11 in the multiplex IHC panel between ILC and IDC/NST in panCK+ tumor cells ( top row) and panCK- stromal cells (bottom row). Data are presented as scatter plots, with each dot representing the mean of percent positive cells for each patient tumor and the solid line indicating the median. Data are analyzed by Mann-Whitney U test. Panels A and B were created in BioRender (Riggins, R. (2026) https://app.biorender.com/illustrations/690fae51cfb5598c629cb341?slideId=c66ed237-665a-43d1-8a8d-0e63a54e1307).

xCT promotes metastasis and therapy resistance in multiple cancers, including breast cancer^19^. As a negative regulator of ferroptosis downstream of xCT, GPX4 expression in cancer and stromal cells is also associated with disease progression and invasiveness^20–22^. GLUD1/2 has recently emerged as a key node connecting glutamate metabolism, xCT function, and ferroptosis^23–25^. However, the contribution of these proteins to ILC biology, prognosis, and clinicopathologic features including tumor characteristics and comorbidities, is not known, despite mounting evidence highlighting the importance of glutamate metabolism in this breast cancer subtype.

## Methods

### Use and Re-Use of Publicly Available and Previously Published Data

mRNA expression data from invasive lobular breast cancer (ILC), mixed ductal-lobular breast cancer (mDLC), and breast cancer of no special type (NST) from The Cancer Genome Atlas (TCGA, RRID:SCR_003193) pan-cancer clinical data resource^26^, METABRIC^27^, and the Sweden Cancerome Analysis Network - Breast (SCAN-B)^28,29^ shown in **Figure S1A** were accessed via Gene Expression eXPLORER (GEXPLORER) at https://leeoesterreich.org/resources on 6/4/2024. mRNA expression data and associated clinicopathological data from ILC and IDC/NST from TCGA shown in **Figure 4C** were accessed via cBioPortal^30,31^ (RRID:SCR_014555) on 6/13/2024. mRNA expression data for *GLUD1* in breast cancer cell lines from DepMap release 25Q2 (https://depmap.org/portal) in **Figure S2A** were accessed on 8/27/2025. GLUD1 peptide counts from ER rapid immunoprecipitation mass spectrometry of endogenous proteins (RIME) in ILC cell lines shown in **Figure 6B** were graphed from Supplemental File 6 of Sottnik et al^32^.

### Primary Breast Cancer Cohorts and Tissue Microarray (TMA) Construction

The ER+ IDC/NST cohort and TMA construction has been previously described^33^. For the ILC cohort, we similarly constructed a TMA with ≥2 cores per patient from n=72 patients with primary ER+ ILC (Inclusion criteria: female, ≥ 10% ERα positivity, PR+/−, HER2 amplification negative, and had surgery for primary breast cancer at MedStar Georgetown University Hospital). Exclusion criteria were patients diagnosed with carcinoma *in situ* only, known *BRCA* or other familial mutation carriers, and evidence of neoadjuvant therapy. All patients were consented through the Histopathology and Tissue Shared Resource, (HTSR), the Survey, Recruitment, and Biospecimen Shared Resource (SRBSR), or IndivuMed under the following respective Georgetown University IRB protocols: 1992-048, Pr0000007, and 2007-345. Demographic, clinical, and pathological data, vital status, and follow-up time (vital status and follow-up time updated July 2021) are shown in **Table 1**.

**Table 1.**
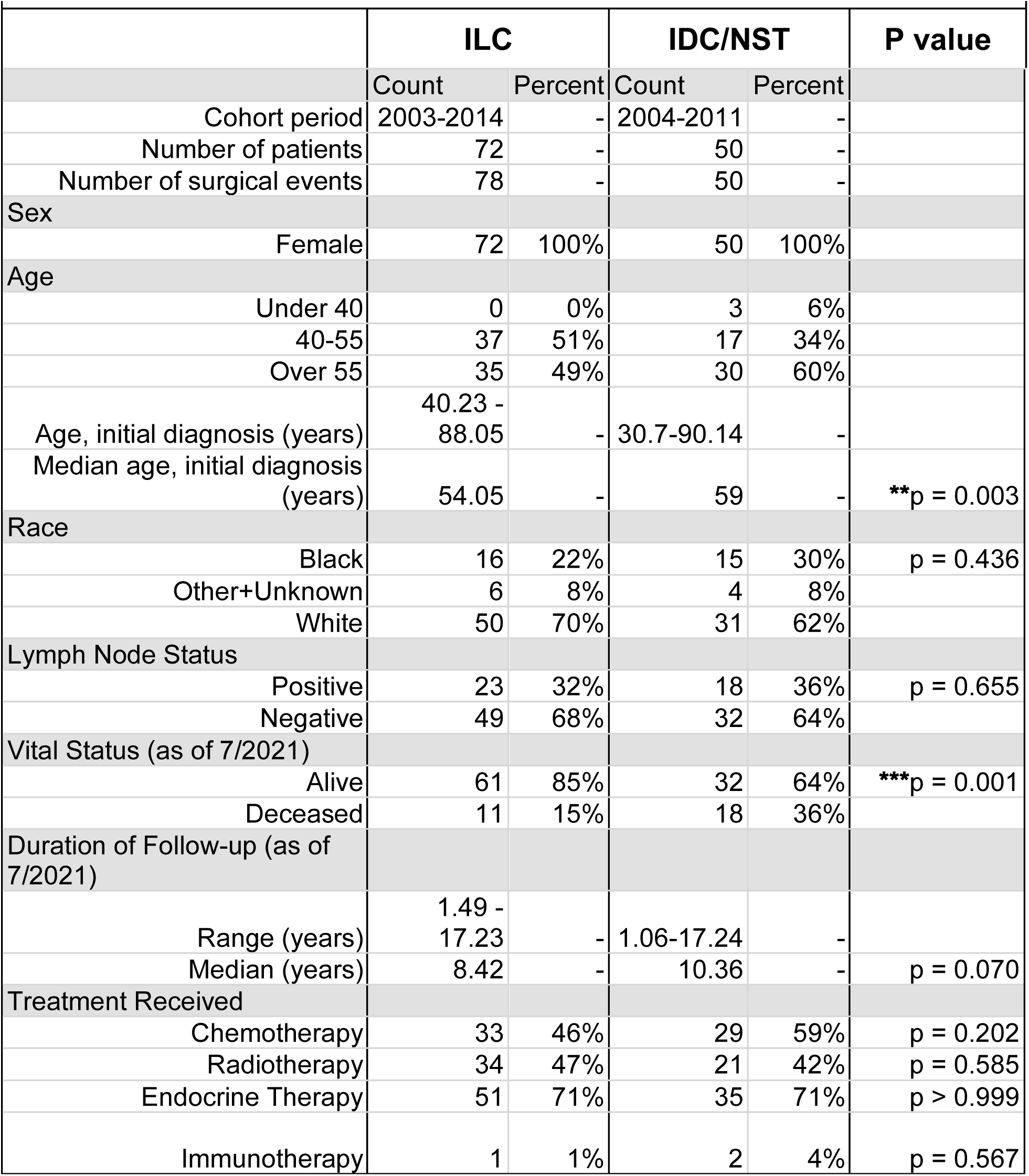
Clinical and pathological characteristics of TMA cohorts.

|  | ILC |  | IDC/NST |  | P value |
| --- | --- | --- | --- | --- | --- |
|  | Count | Percent | Count | Percent |  |
| Cohort period | 2003-2014 | - | 2004-2011 | - |  |
| Number of patients | 72 | - | 50 | - |  |
| Number of surgical events | 78 | - | 50 | - |  |
| Sex |  |  |  |  |  |
| Female | 72 | 100% | 50 | 100% |  |
| Age |  |  |  |  |  |
| Under 40 | 0 | 0% | 3 | 6% |  |
| 40-55 | 37 | 51% | 17 | 34% |  |
| Over 55 | 35 | 49% | 30 | 60% |  |
| Age, initial diagnosis (years) | 40.23 - 88.05 | - | 30.7-90.14 | - |  |
| Median age, initial diagnosis (years) | 54.05 | - | 59 | - | **p = 0.003 |
| Race |  |  |  |  |  |
| Black | 16 | 22% | 15 | 30% | p = 0.436 |
| Other+Unknown | 6 | 8% | 4 | 8% |  |
| White | 50 | 70% | 31 | 62% |  |
| Lymph Node Status |  |  |  |  |  |
| Positive | 23 | 32% | 18 | 36% | p = 0.655 |
| Negative | 49 | 68% | 32 | 64% |  |
| Vital Status (as of 7/2021) |  |  |  |  |  |
| Alive | 61 | 85% | 32 | 64% | ***p = 0.001 |
| Deceased | 11 | 15% | 18 | 36% |  |
| Duration of Follow-up (as of 7/2021) |  |  |  |  |  |
| Range (years) | 1.49 - 17.23 | - | 1.06-17.24 | - |  |
| Median (years) | 8.42 | - | 10.36 | - | p = 0.070 |
| Treatment Received |  |  |  |  |  |
| Chemotherapy | 33 | 46% | 29 | 59% | p = 0.202 |
| Radiotherapy | 34 | 47% | 21 | 42% | p = 0.585 |
| Endocrine Therapy | 51 | 71% | 35 | 71% | p > 0.999 |
| Immunotherapy | 1 | 1% | 2 | 4% | p = 0.567 |

### OPAL Multiplex IHC and Image Processing

Duplicate sections of each TMA, taken from different depths, were subjected to multiplex IHC conducted on the Vectra3 multispectral imaging platform (Akoya Biosciences, Marlborough, MA) using OPAL chemistry as previously described^14^. Antibody/OPAL pairing and staining order was empirically determined by several criteria, including spectrally separating co-localized markers and separating spectrally adjacent dyes. We first performed singleplex IHC with the chosen antibody/OPAL dye pair to optimize signal intensity values and gauge proper cellular expression, followed by optimizing the entire multiplex assay. Primary antibodies in the panel, OPAL pairing, and other methodological details are shown in **Supplemental Table 1**. Image scanning, spectral unmixing, tissue/cell segmentation, phenotyping, and analysis on the Vectra 3.0 Automated Quantitative Pathology Imaging System using Phenochart and inForm 2.4.6 (PerkinElmer/Akoya) was conducted as previously described^14^. Data (percent cell positivity) from replicate cores and duplicate TMA sections for each tumor were averaged.

### Cell Culture and Reagents

Cell lines were maintained at 37°C in a humidified incubator with 95% air and 5% carbon dioxide (CO_2_). Cell lines were authenticated by short tandem repeat (STR) profiling, and regularly tested to ensure they remained free of *Mycoplasma spp*. contamination, by the Tissue Culture and Biobanking Shared Resource (TCBSR). The following ER+ ILC cell lines were cultured as previously described^14,34,35^: SUM44PE (RRID:CVCL_3424) and its tamoxifen-resistant derivative LCCTam; and MDA-MB-134VI (RRID:CVCL_0617) and its long-term estrogen-deprived (LTED) derivatives LTED-A, -B, -D, and -E. General cell culture supplements and reagents were purchased from ThermoFisher (Grand Island, NY) or Sigma Aldrich (St. Louis, MO). The glutamate dehydrogenase inhibitor R162 (Cat# 538098) and Poly-L-lysine hydrobromide (PLL; Cat# P6282) were purchased from Sigma Aldrich.

### Immunoblotting

Assays were conducted as previously described^14^ and whole-cell lysates were probed with the following primary antibodies: SLC3A2 (1:1000, Santa Cruz Biotechnology Cat# sc-376815, RRID:AB_2938854); SLC7A11 (1:500, Abcam Cat# ab175186, RRID: AB_2722749); GLUD1/2 (1:1000, Cell Signaling Technology Cat# 12793, RRID:AB_2750880); GPX4 (1:1000, Abcam Cat# ab125066, RRID: AB_10973901); ER alpha (clone D8H8; 1:1000, Cell Signaling Technology Cat# 8644, RRID:AB_2617128); and beta-actin (1:2000, Cell Signaling Technology Cat# 3700, RRID:AB_2242334). ImageJ^36^ was used to perform densitometry analysis.

### Cell Viability Assays

Cell viability was assessed using the trypan blue exclusion assay. MM134 parental and LTED-D cells were seeded at a density of 100,000 cells/well in 24-well plates and incubated for 24 and 48 hours, respectively, before treatment with the indicated concentrations of R162 for 72 hours. After treatment, cells were trypsinized, stained with trypan blue, and live cells were counted using the Countess II Automated Cell Counter (ThermoFisher Scientific).

### Metabolic Imaging of NADH Autofluorescence, Fluorescent Lifetime Imaging (FLIM) Instrumentation, and Phasor Analysis

MM134 and MM134 LTED-D cells were seeded on PLL-coated 22 mm^2^ diameter glass coverslips (VWR, Cat# 48380-046) in 6-well plates at 150,000 cells/well. Forty-eight hours later, cells were treated with DMSO or 10 μM R162 for 12 hours before transferring coverslips to 35 mm^2^ diameter glass bottom dishes (MatTek, Cat# P35GCOL-1.5-14-C) and performing live-cell imaging on the FVMPE-RS multi-photon, laser-scanning microscope (Olympus, Waltham, MA) with deep imaging via emission recovery (DIVER) detector in the Microscopy and Imaging Shared Resource (MISR). Cells were imaged with a 40X water objective (Olympus, Waltham, MA) using a 740 nm laser from an InsightX3 laser. The signal was collected using the DIVER detector in the forward direction that is connected to a FastFLIM acquisition card (ISS, Champaign IL).

The FLIM data acquisition and analysis was carried out using GSlab^37^. The fluorescence lifetime of each pixel in the autofluorescence intensity image was transformed into a phasor plot. These phasor plots were color-coded based on the phase angle, with longer lifetimes appearing in red–yellow hues and shorter lifetimes in blue–cyan (**Figure S2B**).

To isolate cytoplasmic and mitochondrial regions, a histogram-based intensity correction was applied, excluding nuclear areas due to their lower fluorescence intensity. Fractional intensity of free NADH was calculated using a multiplexing algorithm previously described^14,37–41^. Three phasor components were selected for this analysis: free NADH (0.4 ns), bound NADH (3.4 ns), and background (0.0 ns)^42,43^. Following calculation, individual fractional intensity maps were exported both as data files and as binary images, where fractions ranging from 0 to 1 were scaled to a grayscale range of 0–255. Bound and free NADH fraction images were then merged using ImageJ^36^ to visualize metabolic activity distribution. For enhanced visualization, the color scale was adjusted so that bound NADH (purple) was restricted to a range of 140–255, and free NADH (yellow) to a range of 1–35. This adjustment was necessary due to the predominance of bound NADH in these cell types.

### Metabolic Assays

Non-mitochondrial oxygen consumption rate was determined using Seahorse assays. MM134 parental and LTED-D cells were seeded at 80,000 cells/well in PLL–coated XF96 cell culture microplates (Agilent, Cat# 103793-100). After 24 h, cells were treated with DMSO or R162 (10 or 20 μM) for 12 hours. Media was then replaced with Seahorse XF DMEM supplemented with 10 mM glucose, 2 mM glutamine, and 1 mM sodium pyruvate (Agilent, Cat# 103680-100), followed by 1 hour incubation at 37 °C in a non-CO₂ incubator. Metabolic flux measurements were collected under basal conditions and after sequential injection of the Seahorse Mito Stress Test reagents (Agilent, Cat# 103015-100) at final concentrations of oligomycin (1.5 μM), FCCP (2 μM), and rotenone/antimycin A (0.5 μM), using the Seahorse XFe96 Analyzer (Agilent).

Extracellular lactate was measured using the Lactate-Glo Assay kit (Promega, Cat# J5021) per manufacturer’s instructions. MM134 parental and LTED-D cells were seeded at 25,000 cells/well in 96-well plates precoated with PLL. Parental cells were treated 24 hours after seeding, whereas LTED-D cells were treated 72 hours after seeding, with DMSO or R162 at 10 μM or 20 μM. Media samples were collected 12 hours after treatment. For analysis, 3 µL of media was diluted in PBS (1:200 for parental, 1:300 for LTED-D) and samples were mixed 1:1 with the lactate detection reagent and read on a microplate reader (EnSpire, Perkin Elmer).

### Statistical Analyses

GraphPad Prism 10 (Boston, MA, RRID:SCR_002798) was used for statistical analyses. Mann Whitney tests were used to analyze and compare protein expression in lobular vs. ductal tumor (panCK+) and stromal cells (panCK-), and by lymph node-positive vs. negative status. Simple linear regression and Spearman ρ correlation analyses were used to compare protein expression versus tumor size or patient age, and co-expression between proteins in tumor and stroma. Kaplan-Meier analyses and log-rank tests were used for overall survival analysis. Fisher’s exact and Chi-squared tests were used to analyze demographic, clinical and pathological data where appropriate, and the relationship between comorbidity or race and high vs. low (above vs. below the median) protein expression. Unpaired t-tests and one-way analysis of variance (ANOVA) with Dunnett’s multiple comparison tests were used to analyze cell viability, bound/total NADH from FLIM, Seahorse, and extracellular lactate assays, as appropriate. All relevant tests were two-sided and statistical significance was defined as p≤ 0.05. Asterisks denote statistical significance as follows: *, p≤0.05; **, p≤0.01; ***, p≤0.001; ****, p≤0.0001.

## Results

### Study population and cohort comparisons

We constructed a series of TMAs of ER+ ILC (n=72) and IDC/NST (n=50) for staining with our custom multiplex IHC panel (**Figure 1B**). Our cohort is unique in having 34% of patients self-report their race as non-white, including 25% identifying as Black or African American, with equal representation in the ILC and IDC/NST cohorts **(Table 1)**. There has historically been a paucity of data on ILC from non-white patient populations. However, recent studies report that the average annual percent change in ILC incidence has increased across all racial groups^44^, with a higher increase among non-Hispanic Black women than for non-Hispanic white women in the past few decades^45^. Preliminary evidence suggests that Black women with ILC have significantly lower rates of 5-year breast cancer-specific survival and are less likely to undergo surgery, despite clinically presenting with locally advanced disease^46^.

Prior retrospective analyses suggest that ILC occurs primarily in women >60y, which may reflect challenges in detecting this histological subtype. Another distinct feature of our cohort is that patients with ILC were <60y and younger than those with IDC/NST at diagnosis (54 vs 59y, **p=0.003). Likely consistent with younger age and a numerical trend towards shorter duration of follow-up, a significantly greater proportion of ILC patients were alive at last contact (85%, vs 64% for IDC/NST, ***p=0.001). Other clinicopathological features including nodal status and treatment modality were similar across cohorts. Although all tumors met our inclusion criteria for ER+ and HER2 amplification negative, we further assessed ER, progesterone receptor (PR), and HER2 expression by IHC (Ki67 data were not available) from the original pathology reports. HER2 protein expression (IHC 0 vs. 1+ to 3+) was more common in ILC vs. IDC/NST (Fisher’s exact test *p=0.02). In ILC, percent PR+ was higher but not significant (71% vs. 45% for IDC/NST, p=0.07), whereas percent ER+ was lower (83.5% vs. 91% for IDC/NST, *p=0.02), consistent with a prior study that demonstrated differential ER protein turnover in ILC vs. IDC/NST^47^.

### Expression of glutamate-handling proteins in ILC vs. IDC/NST

We used three well-characterized public datasets (TCGA, METABRIC, and SCAN-B)^26–29^ to compare mRNA expression of the four target proteins in our multiplex IHC panel in ILC, IDC/NST, and (where reported) mixed mDLC (**Figure S1A**). *SLC7A11* and *SLC3A2* expression are both significantly greater in IDC/NST compared to ILC in multiple datasets. This may in part be due to the fact that these analyses do not stratify by hormone receptor status, and xCT is often overexpressed in triple negative breast cancer^48^, which is more prevalent in IDC/NST. By contrast, *GPX4* expression is significantly greater in ILC vs. IDC/NST in the METABRIC dataset, while *GLUD1* expression is significantly greater in ILC vs. IDC/NST in both the METABRIC and SCAN-B datasets. Prior studies in models of mammary epithelial cells show GLUD1 expression is elevated under quiescent and slower growth conditions^49^, which is generally consistent with ILC’s more indolent nature.

In our TMAs, we compared the abundance of each target protein expressed, as a percentage of positive cells, in ILC vs. IDC/NST within the tumor (pan-cytokeratin-positive, panCK+ cells) and stromal (panCK- cells) compartments (**Figure 1C**). Consistent with public mRNA data, but despite all tumors being ER+, SLC7A11 and SLC3A2 protein expression in tumor cells is more abundant in IDC/NST (40-45%) compared to ILC (<25%). In our cohort, GLUD1/2 expression in tumor cells is also significantly greater in IDC/NST, though expression in both cohorts is ≥50%, while there is no significant difference for GPX4 (∼50% positive cells in both cohorts). In stromal or non-epithelial cells, SLC3A2 expression is significantly more abundant in IDC/NST, while GPX4 expression is significantly more abundant in ILC. Together these data suggest that while all four glutamate-handling proteins are expressed in ILC across multiple datasets and sample types, the intracellular enzymes GLUD1/2 and GPX4 are generally more prevalent in ILC vs. IDC/NST.

### All four glutamate-handling proteins are each associated with increased tumor size in ILC

We assessed the relationship between the abundance of each protein in the tumor and stromal compartment, and known breast cancer prognostic factors, including lymph node status, age, and tumor size at diagnosis, using linear regression and Spearman ρ. We find no statistically significant relationship between any of these glutamate handling proteins and either lymph node status or age as a continuous variable (data not shown). However, SLC7A11, SLC3A2, GLUD1/2, and GPX4 expression in tumor cells is each positively and significantly associated with increased tumor size in ILC, but not IDC/NST (**Figure 2A, B**). GLUD1/2 and GPX4 expression in the stroma is also significantly correlated with larger tumors in the ILC cohort. Since larger tumor size is linked to shorter overall survival (OS) in breast cancer (e.g.^50^), we tested whether high vs. low (above vs. below median) protein expression for each target is associated with OS in our ILC and IDC/NST cohorts, which have a median follow-up time of 8.4 and 10.4 years, respectively. Neither individual (data not shown) nor combined expression of the four proteins is significantly associated with OS in either cohort (**Figure 2C**). Together these data highlight a specific association between each member of this network of glutamate-handling proteins and increased tumor size in ILC, but not IDC/NST.

**Figure 2.**
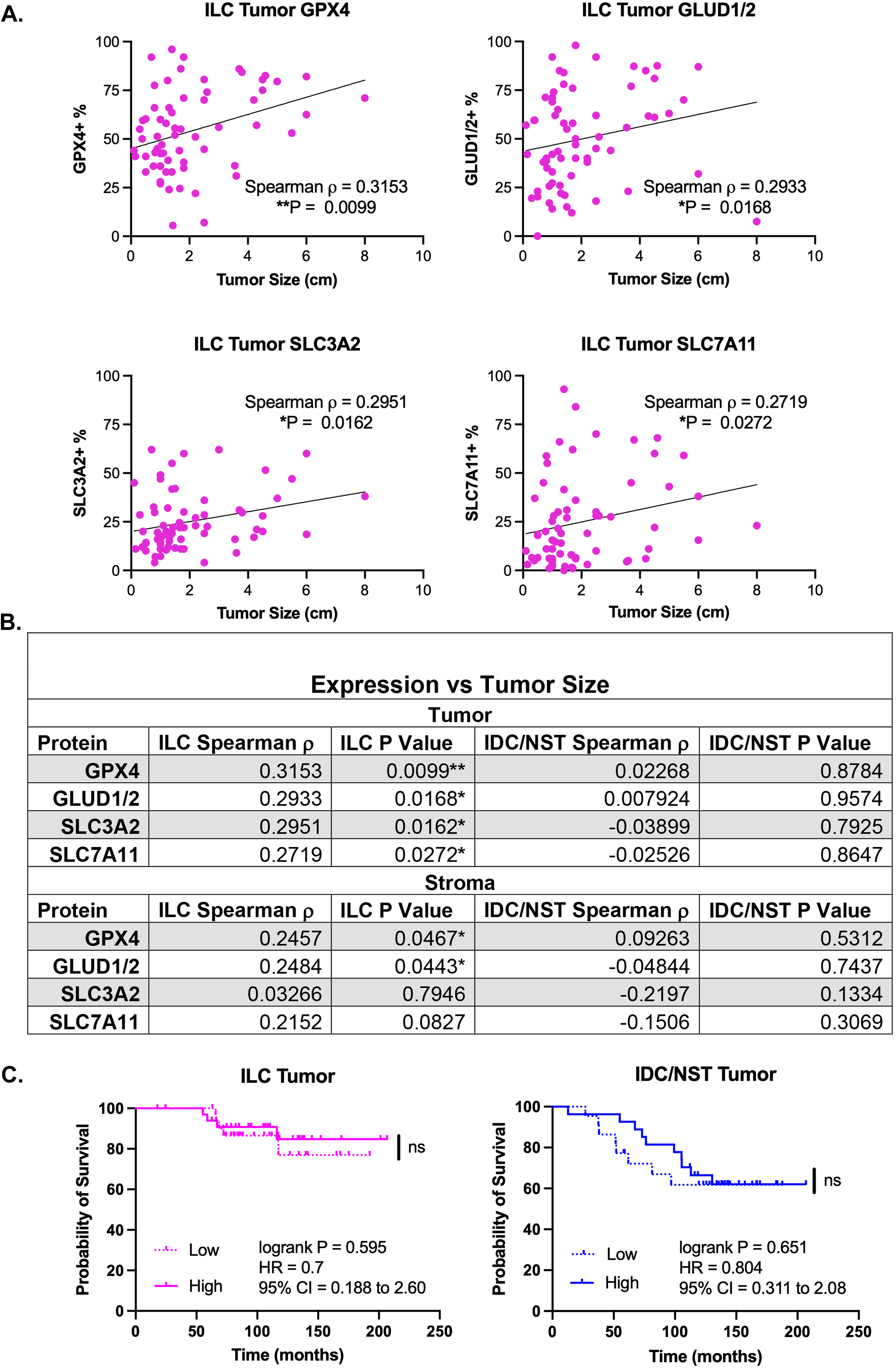
Glutamate-handling proteins are associated with increased tumor size in ILC. **A**, Graphs illustrating the relationship between tumor size (cm) and percent positive cells for tumor expression of GPX4, GLUD1/2, SLC3A2 and SLC7A11 in the ILC cohort. **B**, Spearman correlation coefficient (ρ) and p value for each protein’s relationship to tumor size in panCK+ tumor cells and panCK- stromal cells for the ILC and IDC/NST cohorts. **C**, Kaplan-Meier survival analysis, log-rank p value, hazard ratio (HR), and 95% confidence interval (CI) for high (above median) versus low (below median) tumor expression (percent positive cells) of the combined four proteins in the ILC and IDC/NST cohorts.

### Co-expression of glutamate-handling proteins in ILC

Since SLC3A2, SLC7A11, GPX4, and GLUD1/2 are all functionally related (Figure 1A), we assessed whether they are co-expressed – e.g. do ILC with a high percentage of cells expressing one of the proteins also have a high percentage of cells expressing one or more of the others? There is moderate to strong significant co-expression amongst all four proteins in ILC (**Figure 3A**). For example, in ILC, tumor cell GPX4 abundance is significantly associated with expression of every other protein in both the tumor and the stroma. There is also a significant positive correlation between tumor or stromal GLUD1/2 and ER expression. By contrast, in IDC/NST, tumor cell GPX4 abundance is only significantly positively associated with itself and SLC7A11 in stromal cells, and there is no correlation of any protein with either ER or PR expression (**Figure 3B**). These data show that the known functional relationships between these glutamate-handling proteins translate to increased co-expression in ILC, where all members of this network are significantly associated with increased tumor size.

**Figure 3.**
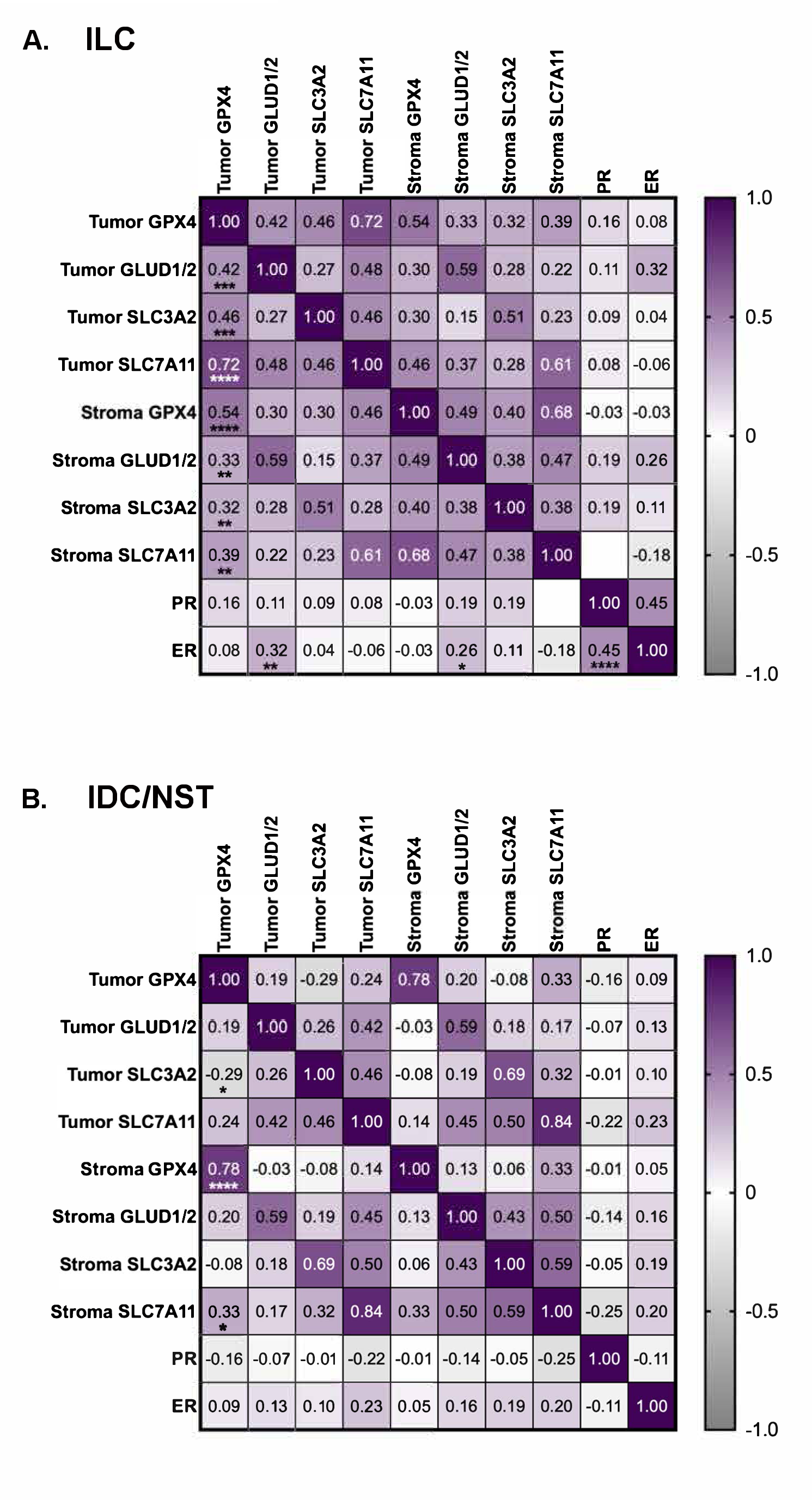
Increased co-expression of glutamate transport proteins and metabolic enzymes in ILC. Heatmaps showing Spearman correlation coefficients (ρ, ranging from +1.0 to -1.0) for each protein’s relationship to one another, and hormone receptor expression (percent positive cells), in the ILC (**A**) and IDC/NST (**B**) cohorts. Asterisks denote statistical significance as defined in the Methods section.

### GLUD1/2, GPX4, and SLC3A2 are strongly associated with increased tumor size in Black women with ILC

Racial disparities in breast cancer outcomes are widely discussed in the context of triple negative breast cancer. However, Parab et al.^51^ recently published that non-Hispanic Black or African-American women are significantly more likely to have aggressive ER+ breast tumors (defined by a high-risk recurrence score on the Oncotype DX panel) when compared to non-Hispanic white women. Van Alsten et al.^52^ show similar data for the PAM50-based risk of recurrence score, wherein young (<50 years of age) Black women with ER+ breast cancer are more likely to have intermediate to high scores as compared to young white women. Consistent with this, Rauscher et al.^53^ report the risk of death from ER+/PR+ breast cancer is more than four times higher for Black women as compared to white women, even after adjustment for tumor stage, grade, and delays in initiation of treatment, while a meta-analysis by Torres et al. shows Black women have a 50% higher relative risk of death from ER+ breast cancer^54^.

Unfortunately, these (and most other) studies of racial disparities in ER+ breast cancer do not report the presence or absence of lobular histology, a point that is complicated by our poor understanding of the true prevalence of ILC in Black women. Analysis of publications that provide such data^55–60^ indicates that the prevalence of ILC in Black women ranges from 5.5% to 16% of breast cancer cohorts – a rate that, in some of these studies, is significantly lower compared to non-Hispanic white women within the same cohort (**Supplemental Table 2**). Consistent with this, Giaquinto et al. report using Surveillance, Epidemiology, and End Results (SEER) data that the age-adjusted incidence rate of ILC for Black women is 11 per 100,000 breast cancer cases vs. 14.7 for non-Hispanic white women^44^. It is important to note that all of these studies may be confounded by variability in diagnostic criteria for ILC in all populations^61,62^.

Nevertheless, with sixteen out of 72 (22%) and fifteen out of 50 (30%) women in our ILC and IDC/NST cohorts, respectively, self-reporting their race as Black or African American (none self-report Hispanic ethnicity), we assessed the relationship between the abundance of each protein in the tumor and stromal compartments with tumor size specifically in Black women. Tumor and stromal cell GLUD1/2, GPX4, and SLC3A2 abundance are each positively and significantly associated with increased tumor size in ILC, but not IDC/NST (**Figure 4A**). For all three proteins, the association with tumor size is markedly stronger in Black women with ILC than in the full ILC cohort (**Figure 4A and 2B**). Although not statistically significant, high individual (data not shown) and combined expression of the four glutamate-handling proteins shows a trend toward worse OS in Black women with ILC (**Figure 4B**).

**Figure 4.**
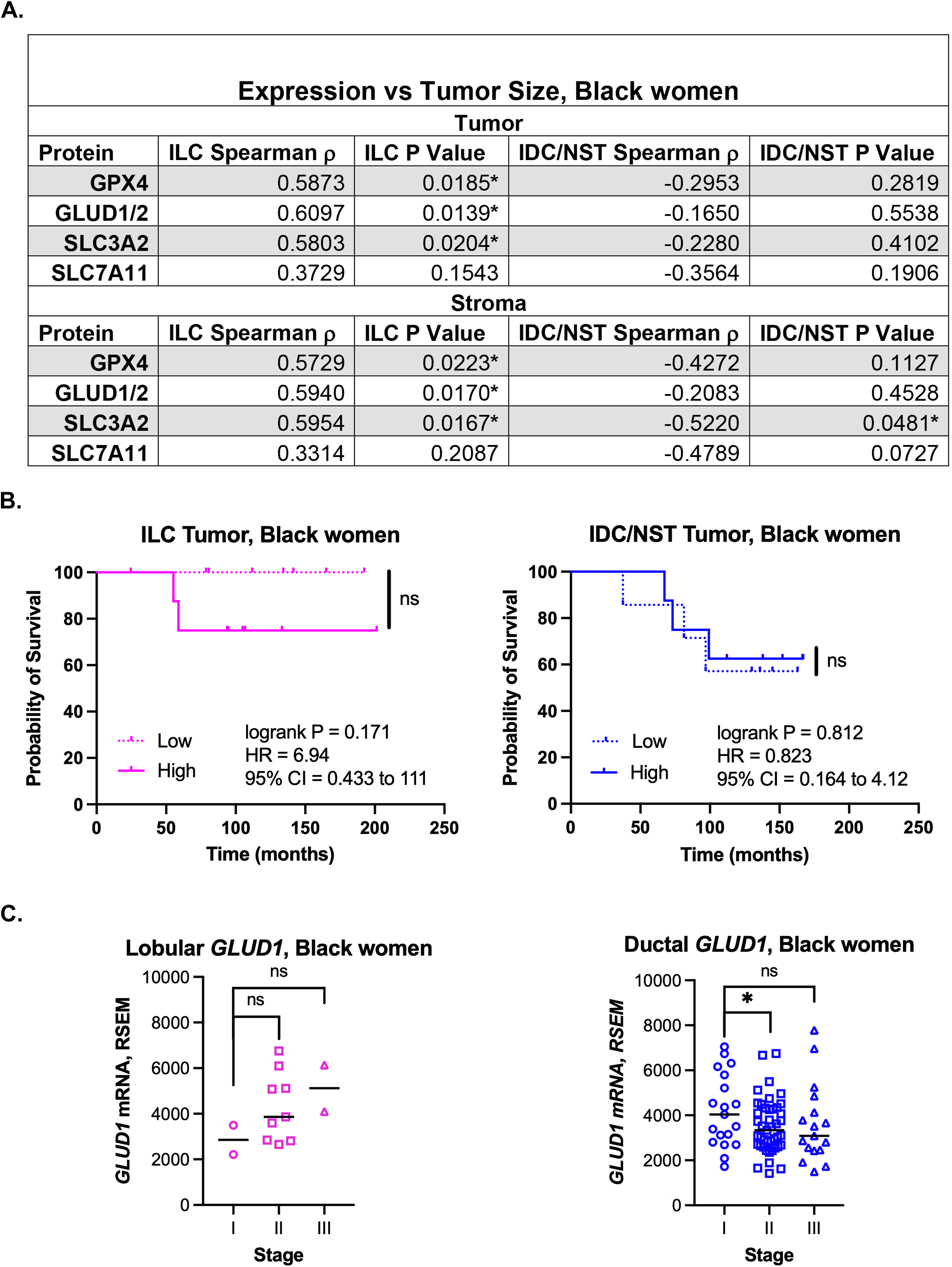
High GLUD1/2 expression is associated with increased tumor size and stage in Black women with ILC. **A**, Spearman correlation coefficient (ρ) and p value for each protein’s relationship to tumor size in panCK+ tumor cells and panCK- stromal cells for Black women in both the ILC and IDC/NST cohorts. **B**, Kaplan-Meier survival analysis, log-rank p value, hazard ratio (HR), and 95% confidence interval (CI) for high (above median) versus low (below median) tumor expression (percent positive cells) of the combined four proteins in Black women with ILC and IDC/NST cohorts. **C**, *GLUD1* mRNA expression (RSEM, RNA-Seq by Expectation-Maximization) by tumor stage from TCGA for Black women with ILC (left) and Luminal A and Luminal B IDC/NST (right) breast cancer. Data are presented as a scatter plot, with the solid line indicating the median. *GLUD1* expression in Stage I vs. Stages II and III by unpaired t-test (*, p<0.05).

Adding to our poor understanding of the ILC biology in Black women is the limited racial diversity in publicly available datasets. METABRIC and SCAN-B report zero Black women with ILC, and current TCGA data report only thirteen (6.5% of the ILC cohort). We first analyzed *GLUD1* mRNA expression in relation to tumor stage in Black women with ILC or with luminal IDC/NST in TCGA (**Figure 4C**), since this enzyme has the strongest correlation with tumor size (Spearman ρ=0.61) and is uniquely associated with ER expression in our ILC cohort (**Figure 3A**). Despite the low sample number, there is a clear, stage-dependent increase in *GLUD1* expression in tumors from Black women with ILC, though not statistically significant (**Figure 4C**). By contrast, there is a significant decrease in *GLUD1* expression in Stage II vs. Stage I luminal IDC/NST in Black women. These stage-dependent trends are not observed for *GPX4* or *SLC3A2* mRNA expression in Black women of either cohort (data not shown). Gene set enrichment analysis (using Enrichr^63–65^) of the top 100 genes positively correlated with *GLUD1* expression in TCGA tumors from Black women with ILC show an enrichment for glutamine and glutamate metabolism pathways (KEGG, q=0.011). Tumors from Black women with ILC also show significant enrichment for the synthesis of arginine (KEGG, q=0.011), N-glycans (KEGG, q=0.046), and uridine diphosphate N-acetylglucosamine (UDP-GlcNAc, Metabolomics Workbench Metabolites, q=0.018). With each of these ILC-enriched pathways independently linked to breast cancer progression and metastatic potential (e.g.^66,67^), this analysis further highlights the potential importance of GLUD1/2 for Black women with ILC.

### Enrichment of GLUD1/2 and SLC3A2 in women with ILC and hypertensive comorbidities

Multiple comorbid conditions like hypertension, obesity and metabolic syndrome are linked to poor outcomes in breast cancer, and there are racial disparities in the prevalence of these comorbidities^68–71^. Abstraction of electronic health records show no difference in the presence or absence of total comorbidities by ICD-9-CM diagnosis codes (details in **Supplemental Table 3**). However, obesity is significantly enriched among patients with IDC/NST, while hypertension is the most prevalent comorbidity in both the ILC and IDC/NST cohorts, with no statistically significant difference between them (**Figure 5A**). Next, we compared comorbidity presence or absence between Black and white women. Significantly more Black women have at least one comorbid condition in both cohorts (**Figure 5B**). Of note, hypertension is significantly enriched among Black women with ILC, but not IDC/NST (**Figure 5C**). Among women with ILC, high expression of GLUD1/2 and SLC3A2 (above median) is enriched in hypertensive patients in both the full and Black cohorts (**Figure 5D** and **5E**), indicating an association between hypertension and elevated expression of glutamate-handling proteins in ILC.

**Figure 5.**
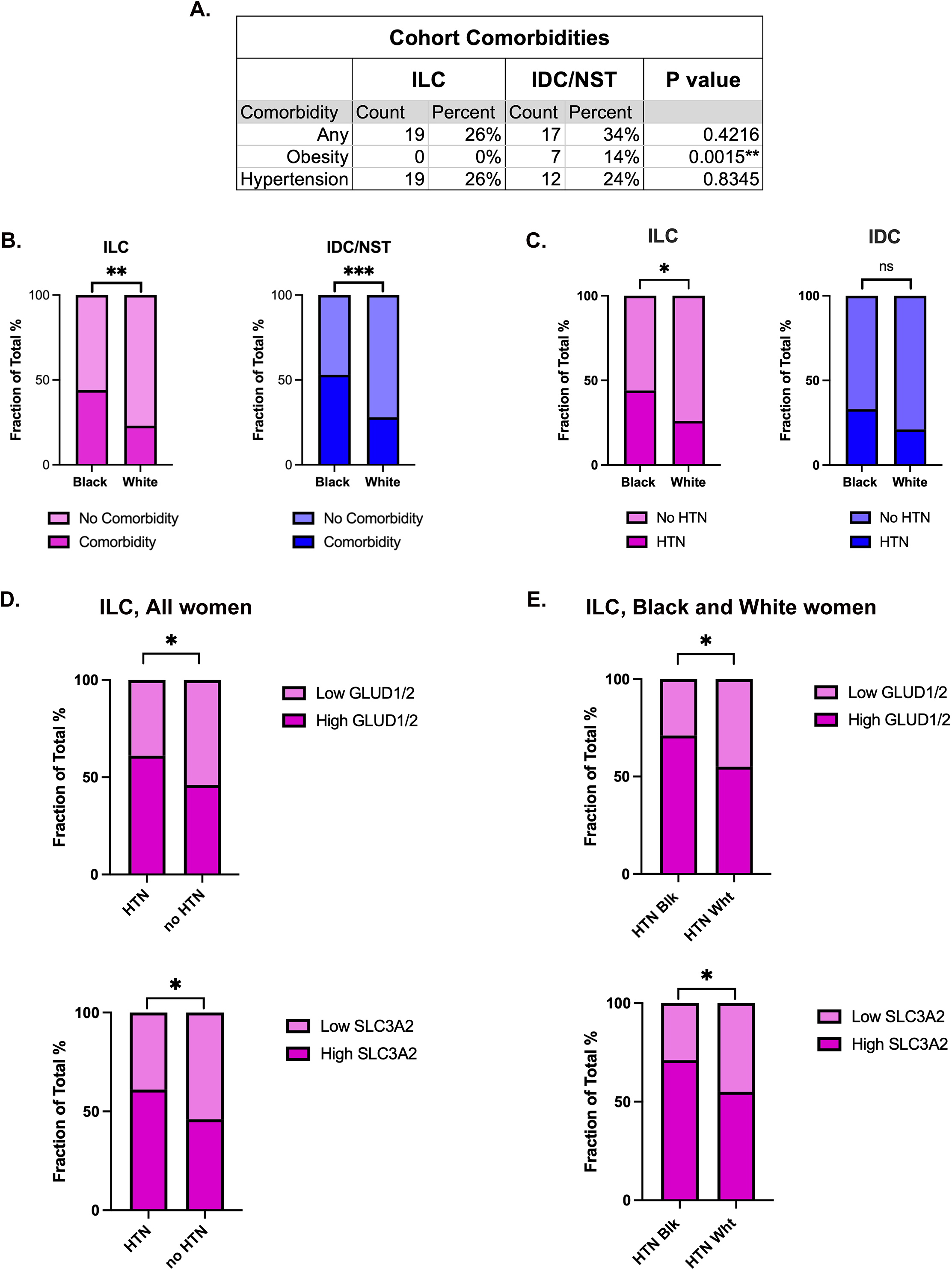
Enrichment of GLUD1/2 and SLC3A2 in tumors from women with hypertensive comorbidities in ILC. **A**, Fisher’s Exact Test of the presence or absence of any comorbidity, or specific comorbidities, in ILC compared to IDC/NST. **B-C**, Fisher’s Exact Test of the presence or absence of any comorbidity (B) or hypertension (HTN; C) in Black women compared to white women across the ILC and IDC/NST cohorts. **D-E**, Fisher’s Exact Test of the proportion of high (above median) and low (below median) GLUD1/2 and SLC3A2 expression in women with versus without HTN in the entire ILC cohort (left), and in hypertensive Black (HTN Blk) vs White (HTN Wht) ILC cohorts (right).

### Pharmacological inhibition of GLUD1 reduces ER protein levels, cell viability and oxidative phosphorylation in ILC cell lines

Breast cancer preclinical models are not racially or ancestrally diverse. Nevertheless, we used two of the best-characterized ER+ ILC cell line models and their endocrine therapy-resistant derivatives^34,35^, to evaluate the expression of the four target proteins by immunoblotting (**Figure 6A**). Tamoxifen-resistant LCCTam cells show modest (<2-fold) upregulation of SLC7A11, GLUD1/2, and GPX4 compared to the parental SUM44 cells. However, three of the four long-term estrogen-deprived cell lines (LTED, mimicking aromatase inhibitor resistance) derived from parental MDA-MB-134VI (MM134) cells exhibit robust, 2- to 6-fold upregulation of GPX4 and GLUD1/2, suggestive of a role for these enzymes in aggressive or more advanced ILC.

**Figure 6.**
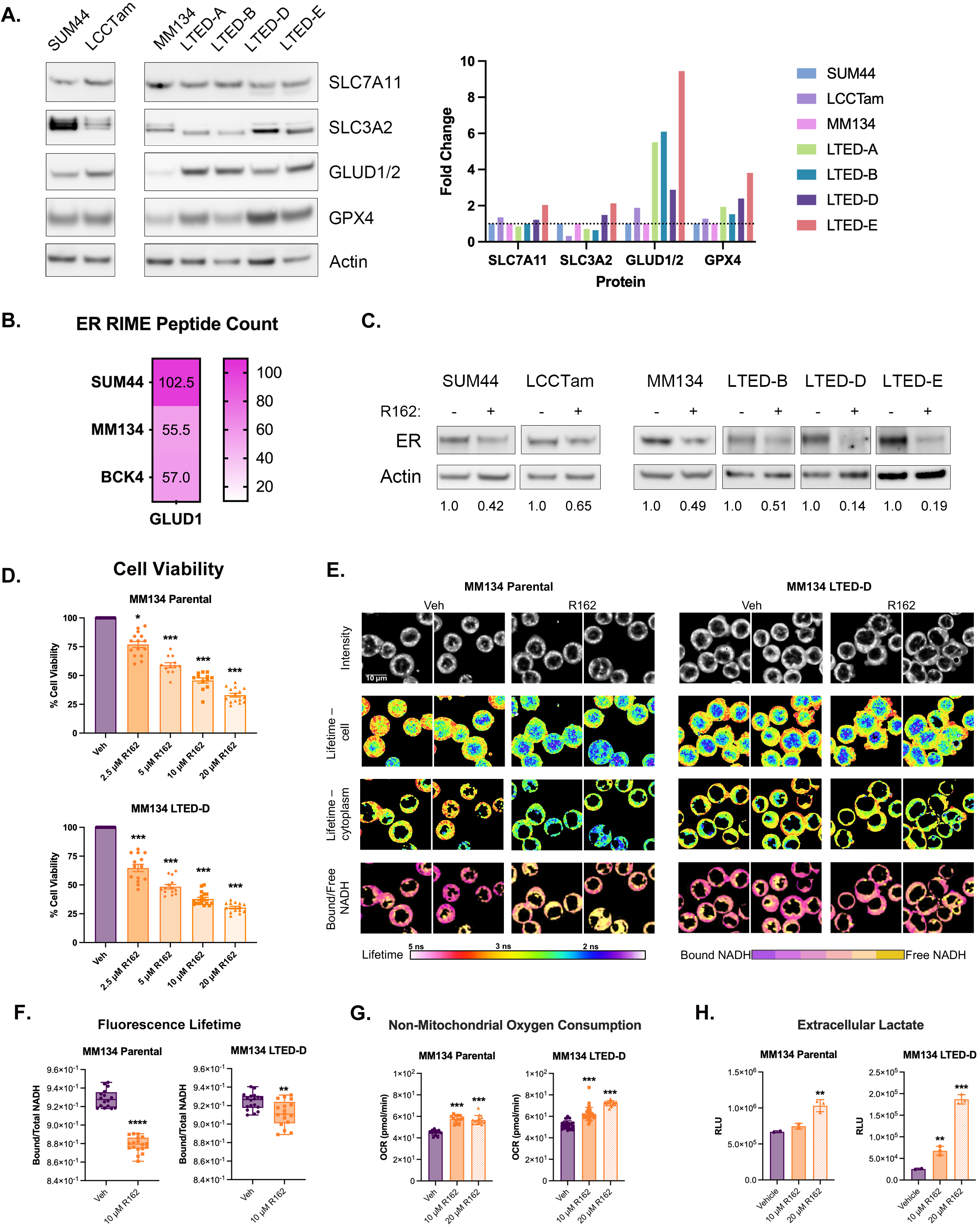
Pharmacological GLUD1 inhibition reduces ER protein levels, cell viability and oxidative phosphorylation in ILC cell lines. **A**, Left, immunoblot analysis of the expression of proteins in ILC breast cancer cell lines. SUM44 and LCCTam, pair of tamoxifen-sensitive and - resistant (respectively) cell lines. MM134 and LTED-A through E, series of parental (MM134) and long-term estrogen-deprived (LTED, mimicking aromatase inhibitor resistance) cell lines. Actin serves as a loading control. Right, densitometry plot showing fold change in expression (target:actin ratio) for each resistant cell line normalized to its parental control. **B**, Heatmap of GLUD1 peptide count from ER RIME assays published by Sottnik et al^32^. Data are presented as the mean peptide count from two biological replicates of the SUM44, MM134, and BCK4 ILC cell lines. **C**, Immunoblot analysis of the expression of ER in ILC breast cancer cell lines cultured in the absence (-) and presence (+) of treatment with 10 μM GLUD1 inhibitor R162 for 18-24 hours. Actin serves as a loading control. ImageJ was used to quantify relative expression in R162 - vs. DMSO-treated cells. **D**. Cell viability after treatment of ILC cells with increasing doses of R162 for 3 days. Data were analyzed from three independent experiments, each with 4-8 technical replicates. **E**, Fluorescence lifetime imaging microscopy (FLIM) of NADH in MM134 parental and MM134 LTED-D cells under vehicle and R162 treatment conditions. The top row displays autofluorescence intensity images in the NADH channel. Row 2 shows phase lifetime–color-coded images, where shorter lifetimes (blue–cyan) are predominantly nuclear and longer lifetimes (yellow–red) are cytoplasmic. Row 3 presents histogram-corrected images highlighting the distribution of lifetimes in only the cytoplasm, with a notable shift toward shorter lifetimes (blue–green) in R162-treated parental cells. Row 4 shows color-coded maps of bound/free NADH fractions, indicating increased free NADH (yellow) and decreased bound NADH (purple) upon R162 treatment, especially in parental cells. Scale bar = 10 μm. **F**, Quantification of bound/total NADH fractional intensity ratios derived from FLIM images in A. Data are presented as a box and scatter plot, with each point representing a single image (16-18 per treatment group) and the solid line indicating the median. Data are analyzed by unpaired t-test. **G**, Non-mitochondrial oxygen consumption in ILC cells treated with vehicle or R162. Data were analyzed from two independent passages for parental cells and four independent passages for LTED-D cells, each with 6-8 technical replicates. **H**, Extracellular lactate production in culture media from ILC cells treated with vehicle or R162. Data were analyzed from two independent experiments, each with three technical repeats, and a representative graph from one experiment is shown. Data in D, G and H are presented as mean ± SEM and analyzed by one-way ANOVA with Dunnett’s multiple comparison test. Asterisks denote statistical significance as defined in the Methods section.

Given this pattern, we next focused on GLUD1 to extend our novel observations that GLUD1/2 expression is: 1) markedly increased in endocrine therapy-resistant variants of ER+ ILC cells (**Figure 6A**); 2) significantly associated with increased tumor size and ER expression in our ILC patient cohort (**Figures 2-4**); and **3**) upregulated with increasing stage in tumors from Black women with ILC in TCGA (**Figure 4C**). Among breast cancer cell lines profiled by DepMap^72^, ER+ ILC and ILC-like cell lines (as defined in^73^) rank among the highest for *GLUD1* mRNA expression (**Figure S2A**). Given its dual roles in TCA cycle anaplerosis and ammonia recycling, GLUD1/2 is predominantly localized in the mitochondria. However, a sub-population of GLUD1 has been reported in the nucleus, where it is involved in chromatin regulation and gene transcription in a catalytic activity-dependent manner^74,75^. Proteomic profiling of the nuclear ER interactome with ER RIME (rapid immunoprecipitation mass spectrometry of endogenous proteins) by Sottnik et al.^32^ identified GLUD1 as one of the top ILC-specific ER-associated proteins (**Figure 6B**) that was not identified in IDC/NST cells^76^. To examine the potential role of GLUD1 in regulating ER expression, we treated four ER+ ILC cell lines with the GLUD1 inhibitor R162. Short-term treatment with R162 reduces ER expression by 35-86% in all cell lines tested (**Figure 6C**), including endocrine therapy-resistant cell lines.

To assess whether GLUD1 can be a useful target for therapeutic intervention in ILC, we performed cell viability assays and show that R162 significantly reduces cell viability of parental MM134 and its endocrine therapy-resistant derivative LTED-D cells in a dose-dependent manner (**Figure 6D**). Previous studies showed that R162 treatment attenuates GLUD1’s pro-tumorigenic functions in multiple malignancies by reducing alpha-ketoglutarate levels and inducing cell death by blocking mitochondrial oxidative phosphorylation (OXPHOS) and/or increasing reactive oxygen species (ROS)^17,77–80^. To evaluate potential metabolic reprogramming in response to GLUD1 inhibition in ILC cells, we used an innovative metabolic imaging approach that combines fluorescence lifetime imaging (FLIM) with phasor analysis (**Figure 6E, S2B**). FLIM leverages autofluorescence of the reduced form of nicotinamide adenine dinucleotide (NADH) and permits separation of protein-bound vs. free NADH, which correlates with OXPHOS vs. glycolytic metabolism, respectively^14,39,81^. In both parental MM134 and LTED-D cells, R162 significantly reduces the proportion of protein-bound NADH, suggestive of reduced OXPHOS (**Figure 6F**). To confirm these results, we performed conventional metabolic assays. R162 concomitantly increases non-mitochondrial oxygen consumption and extracellular lactate production (**Figure 6G, 6H**), each of which can be a measure of reduced OXPHOS and increased ROS associated with mitochondrial dysfunction^82–84^. Altogether, GLUD1 pharmacological inhibition suppresses cell viability, reduces ER protein expression, and reprograms metabolism in multiple ILC cell lines, providing preliminary evidence that inhibition of GLUD1 may offer therapeutic benefit in ILC.

## Discussion

ILC has emerged as a breast cancer histologic subtype that is more reliant on amino acid and lipid metabolism than glucose metabolism. Our prior studies specifically highlight the importance of glutamate signaling in ILC^13,14^. Here, we provide evidence for a relationship between a focused network of four glutamate-handling proteins (SLC3A2, SLC7A11, GPX4, and GLUD1/2) and increased tumor size in primary ILC, but not primary IDC/NST. GLUD1/2, GPX4, and SLC3A2 expression in tumor and stroma show a markedly stronger relationship with increased tumor size in Black women with ILC. We further demonstrate that GLUD1 and GPX4 are upregulated in ILC endocrine therapy-resistant preclinical models, and that pharmacological inhibition of GLUD1 reduces cell viability, in part by reducing ER protein levels and OXPHOS.

All four glutamate metabolism and transport proteins measured here each have a significant positive relationship with larger tumor size, and with each other, in ILC. Their high degree of co-expression results in multicollinearity that confounds multiple linear regression models, therefore, we do not report the results of these analyses. The strong association of individual and combined protein expression with tumor size, but lack of a significant association with overall survival in our cohort, could be explained by limited statistical power to detect survival differences in a cohort of this size. Larger studies are needed to better assess these relationships. Future comprehensive *in vitro* and *in vivo* preclinical studies will also be necessary to determine if targeting one protein impacts expression or function of the others in ILC.

This interdependency is particularly relevant for assessing the role of SLC7A11 in ILC, since it not only heterodimerizes with SLC3A2, but this heterodimer acts directly upstream of the ferroptosis suppressor GPX4 (discussed further below). ER can transcriptionally upregulate SLC7A11, which is further elevated in endocrine therapy-resistant ER+ breast cancer^85^. However, these studies by Cao et al. were conducted in preclinical models of IDC/NST, and future studies will be needed to specifically address the role of SLC7A11 in ILC.

In addition to its association with tumor size, high expression of SLC3A2 is enriched among hypertensive women in our full and Black ILC cohort. SLC3A2 heterodimerizes with SLC7A8 (LAT2), and genetic variation in these genes has been linked to hypertension risk, supporting a role for altered amino acid transport in regulating renal dopamine synthesis, sodium handling, and blood pressure^86^. Upregulation of SLC3A2 is also associated with poor prognosis in multiple cancers^21,87,88^, including certain highly proliferative, c-Myc-driven breast cancers^22^. Recent studies show SLC3A2 is required for the growth of tamoxifen-resistant ER+ breast cancer cells^89^. Importantly, SLC3A2 is an obligate heterodimeric partner for multiple L-type transporters beyond SLC7A11 and SLC7A8, positioning this protein as a crucial node in a broader amino acid transport network. For example, heterodimerization of SLC3A2 with SLC7A5 (LAT1) is essential for pro-survival amino acid transport in multiple preclinical cancer models^90^. Independent of amino acid transport, SLC3A2 acts as a beta1 and beta3 integrin co-receptor that is implicated in cell adhesion and mechanotransduction^91^, both of which are compromised in ILC as a result of *CDH1* mutational inactivation. The relative contribution of SLC3A2 to amino acid transport-dependent and -independent functions in ILC remains to be determined.

GPX4 plays a key role in detoxifying lipid peroxides that accumulate in cells, thereby preventing ferroptosis^92,93^. Recent work by Hu et al. shows that endocrine therapy can sensitize ER+ IDC/NST cell lines to ferroptosis induction by GPX4 inhibitors^94^, while in tamoxifen-resistant IDC/NST cell lines, activation of the RelB arm of the NF-kB pathway inhibits ferroptosis by upregulating GPX4^95^. It is not known whether RelB is a transcriptional regulator of GPX4 in ILC, but we do see an increase in GPX4 expression in tamoxifen-resistant and LTED ILC cells (**Figure 6A**). Of note, GPX4 sits at the intersection of multiple pathways that confer heightened sensitivity to ferroptosis. Cell-cell interactions mediated by E-cadherin suppress ferroptosis due to activation of the Hippo pathway, and loss or inhibition of E-cadherin leads to increased vulnerability to ferroptosis^92,96^. Cell density also influences ferroptosis sensitivity in the presence of wild type E-cadherin. Cells cultured at low density with few cell-cell contacts are more susceptible to cell death caused by ferroptosis via GPX4 inhibition than those cultured at high density^93^. In the stroma, ferroptosis and GPX4 are regulated by metabolic pathways that influence tumor crosstalk with the potential to alter cancer progression and response to treatment (discussed in ^97,98^). With E-cadherin loss via *CDH1* mutational inactivation and a discohesive growth pattern with few cell-cell contacts both being hallmarks of ILC that contribute to ferroptosis sensitivity, our data strongly implicate GPX4 as an attractive target for therapeutic intervention in ILC that requires further exploration.

GLUD1/2 protein expression in both the tumor and stromal compartments of ILC is strongly correlated with increased tumor size in Black women in our cohort, while *GLUD1* mRNA increases with tumor stage in ILC from Black women in TCGA (**Figure 4**). By contrast, there is an inverse or negative relationship between GLUD1/2 protein expression and tumor size, and *GLUD1* and tumor stage in luminal IDC/NST, in our cohort and in TCGA data, respectively. Our findings in ILC contradict the observations of Craze et al.^99^, who showed that *GLUD1* mRNA decreases with tumor grade, while both mRNA and protein are associated with better breast cancer-specific survival. Their study included ILC cases (∼8%), but the analyses did not separate or compare ILC with IDC; we therefore posit that the reduction of GLUD1 seen with increasing tumor grade in their study stems from the IDC/NST cases. Also in that work, the IDC/NST category explicitly included mDLC, which we now appreciate as highly heterogeneous and perhaps a distinct molecular entity^100^. The strong association of GLUD1/2 with tumor size and stage in Black women, but lack of a significant association with overall survival in our cohort, could be explained by limited statistical power to detect survival differences, given the relatively small subset of Black women with ILC in our cohort. Larger studies will be needed to better assess these relationships.

Findings from our preclinical studies further support the conclusion that GLUD1/2 expression may confer poor prognosis in ILC. GLUD1/2 is increased in tamoxifen-resistant and LTED ILC cells, has novel interaction with nuclear ER in ILC cell lines (as demonstrated by RIME^32^), and pharmacological inhibition of GLUD1 markedly reduces ER protein expression in multiple ILC cell lines (**Figures 6A-6C**). Placed in the context of current literature demonstrating ER’s novel transcriptional regulatory functions and coregulators in ILC^32,35,101,102^, and the role of a nuclear GLUD1 sub-population in chromatin remodeling and gene transcription^74,75^, we propose that GLUD1:ER interactions may confer ILC-specific ER activities that could be targeted by GLUD1 inhibition, alone or in combination with endocrine therapies. Accordingly, testing whether GLUD1 inhibition synergizes with endocrine therapy and/or reverses endocrine resistance in ILC is warranted. It is important to note that while we show through biochemical and advanced microscopy approaches that pharmacological inhibition of GLUD1 in ILC cells leads to a decrease in OXPHOS, further studies are needed to distinguish these well-studied mitochondrial functions of GLUD1 from its still-emerging roles in the nucleus. GLUD1 inhibition has also been reported to induce ferroptosis in other cancers^17^. The simultaneous R162-mediated increase in extracellular lactate and non-mitochondrial oxygen consumption we observe in ILC cells is suggestive of oxidative stress, which can lead to ferroptosis^103^. Therefore, like GPX4 (discussed above), we posit that targeting GLUD1 may also represent a therapeutic opportunity to capitalize on ILC’s hypothesized increased sensitivity to ferroptosis.

There are several limitations to consider in the interpretation of this study. This is a single-institution, relatively small cohort, despite better proportional representation of Black or African American women than many other published studies of ILC. Self-reported race is an important social construct, but ancestry informative marker-guided analyses will be integral to our follow-up studies, as will access to more comprehensive data on comorbidities, treatments received, and systemic and social determinants of health that are potential mediators of breast cancer disparities. This latter point is particularly relevant because several of the glutamate-handling proteins studied here have mechanistic connections to biological readouts for stress (such as allostatic load) that is disproportionately experienced by minoritized or marginalized communities, and/or associated with higher breast cancer risk or poorer outcomes^104–107^. To our knowledge just two studies have specifically examined systemic and social determinants of health in ILC. Kaur et al.^108^ report that lower socioeconomic status is associated with larger tumor size, higher tumor grade, and an increased risk of death in ILC, while the increased incidence of ILC reported by Quinn et al.^45^ is particularly acute in areas with the lowest attainment of high school education (<10%). Our ongoing studies seek to integrate these systemic and social determinants of health with genomic and transcriptomic patterns in large, diverse, multi-institutional ILC cohorts.

In conclusion, our novel findings implicate an important role for glutamate handling in ILC and suggest GLUD1 and potentially other glutamate-handling proteins as candidate targets for therapeutic intervention, warranting further investigation in diverse preclinical and translational studies.

## Author Contributions

Conception and design: TAY, SB, RBR

Provision of study materials: AMR, KC, BTH

Collection and assembly of data: TAY, SB, TCA, EAP, ATR, NH, SP, AIF, DM, AOO, SR, JLS, MJS

Data analysis and interpretation: TAY, SB, TCA, EAP, ATR, AOO, SR, JLS, MJS, ZME Manuscript writing: All authors.

Final approval of manuscript: All authors.

Accountable for all aspects of the work: All authors.

## Ethical Approvals

All individuals whose breast tumor samples were included on the TMAs were research-consented through the Histopathology and Tissue Shared Resource, (HTSR), the Survey, Recruitment, and Biospecimen Shared Resource (SRBSR), and/or IndivuMed under the following Georgetown University IRB protocols: 1992-048, Pr0000007, and 2007-345.

## Preprints, Prior Publication, and Author Self-Archiving

The Author’s Original Version of this manuscript has been preprinted at bioRxiv. This does not infringe any subsequent copyright or license agreement.

## Acknowledgements

We are grateful to the women who so generously consented to donate their breast tumors for research. We wish to thank members of the Riggins Lab (especially Stanley Tam, Angela Appiah-Kubi, Alicia Nash, and Melody Kabbai), members of NR IMPACT, the Hackensack Meridian Health-John Theurer Cancer Center Community Advisory Council, as well as Anju Duttargi, Raneen Rahhal, and Roberto D’Angelo-Cosme for their insights and helpful suggestions. This work was financially supported by: Department of Defense Breast Cancer Research Program Award W81XWH-17-1-0615, NIH/NCI Cancer Center Support Grant (CCSG) Administrative Supplement P30 CA051008-31S1, and BellRinger pilot funds from Georgetown Lombardi to RBR; American Cancer Society Institutional Research Grant (ACS IRG) pilot award from IRG- 23-1156148-27-IRG to SB; CCSG Developmental Funds from P30 CA051008 and NIH/NIGMS R35 GM154815 to SR; and NIH/NCI R00 CA193734 and R01 CA251621 to MJS. Fellowship funding for TAY was provided by the Tumor Biology Training Grant T32 CA009686. TAY also received a travel award to present a portion of this work at the 2023 Endocrine Society Annual Meeting. Fellowship funding for TCA and EAP was provided by ACS Center for Innovation in Cancer Research Training awards IRG-17-177-23-IRG and DICR INTR-23-1253711-01-DICR INTR. TCA also received a travel award to present a portion of this work at the 2023 Annual Biomedical Research Conference for Minoritized Scientists (ABRCMS). SP received a Georgetown Undergraduate Research Opportunities Program (GUROP) Summer Fellowship. DM received a Summer Mentored Undergraduate Research Fellowship (SMURF) award from Georgetown University, and support from a 2024-2025 Goldwater Scholar award. Technical services were provided by the TCBSR, HTSR, SRBSR, MISR, and FCSR, as described in the Methods section, which are supported in part by P30 CA051008. The content of this article is the sole responsibility of the authors and does not represent the official view of the DoD, NIH, ACS, or the Goldwater Scholarship Foundation.

## Abbreviations

[^18^F]FDG-PET: 18F-fluorodeoxyglucose positron emission tomography
Asp: aspartic acid
Cys: cyst(e)ine
DIVER: Deep Imaging Via Emission Recovery
ER+: Estrogen Receptor Positive
FLIM: Fluorescent Lifetime Imaging
GEXPLORER: Gene Expression eXPLORER
Gln: glutamine
Glu: glutamate
GLUD1/2 or GDH1/2: Glutamate Dehydrogenase 1/2
Gly: glycine
GPX4: Glutathione Peroxidase 4
GSH: glutathione
GSSG: glutathione disulfide
HTN: hypertension
IDC/NST: Invasive Ductal, or Breast Cancer of No Special Type
IHC: Immunohistochemistry
ILC: Invasive Lobular Breast Cancer
LCCTam: Tamoxifen-resistant Derivative of SUM44PE
Leu: leucine
LTED: Long-term Estrogen-deprived Derivative of MDA-MB-134VI
mDLC: Mixed Ductal-lobular Breast Cancer
mGluRs: Metabotropic Glutamate Receptors
OXPHOS: Oxidative Phosphorylation
panCK: Pan-cytokeratin
Pro: proline
R162: GLUD1 inhibitor
RIME: Rapid Immunoprecipitation Mass Spectrometry of Endogenous Proteins
ROS: Reactive Oxygen Species
SCAN-B: Sweden Cancerome Analysis Network – Breast
SEER: Surveillance, Epidemiology, and End Results
SLC: solute carrier
TCA: tricarboxylic acid
TCGA: The Cancer Genome Atlas
TMA: Tissue Microarray.

## Supplemental Figure Legends

**Figure S1.**
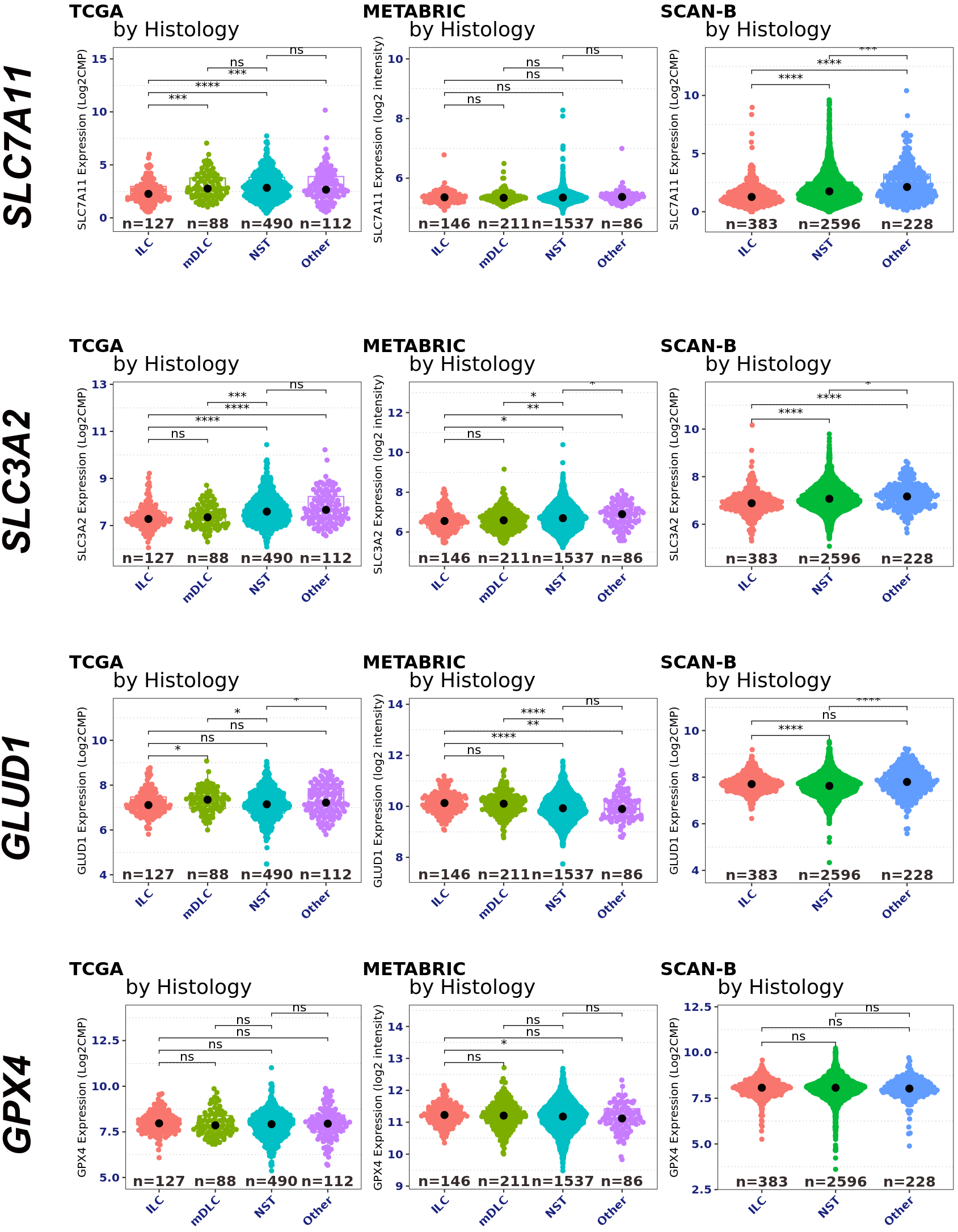
Expression of glutamate-handling proteins in publicly available breast cancer datasets. Violin plots of mRNA expression data from invasive lobular breast cancer (ILC), mixed ductal-lobular breast cancer (mDLC), and breast cancer of no special type (NST) for each target in the multiplex IHC panel from TCGA, METABRIC, and SCAN-B. Data were analyzed and graphed using Gene Expression eXPLORER (GEXPLORER) at https://leeoesterreich.org/resources.

**Figure S2.**
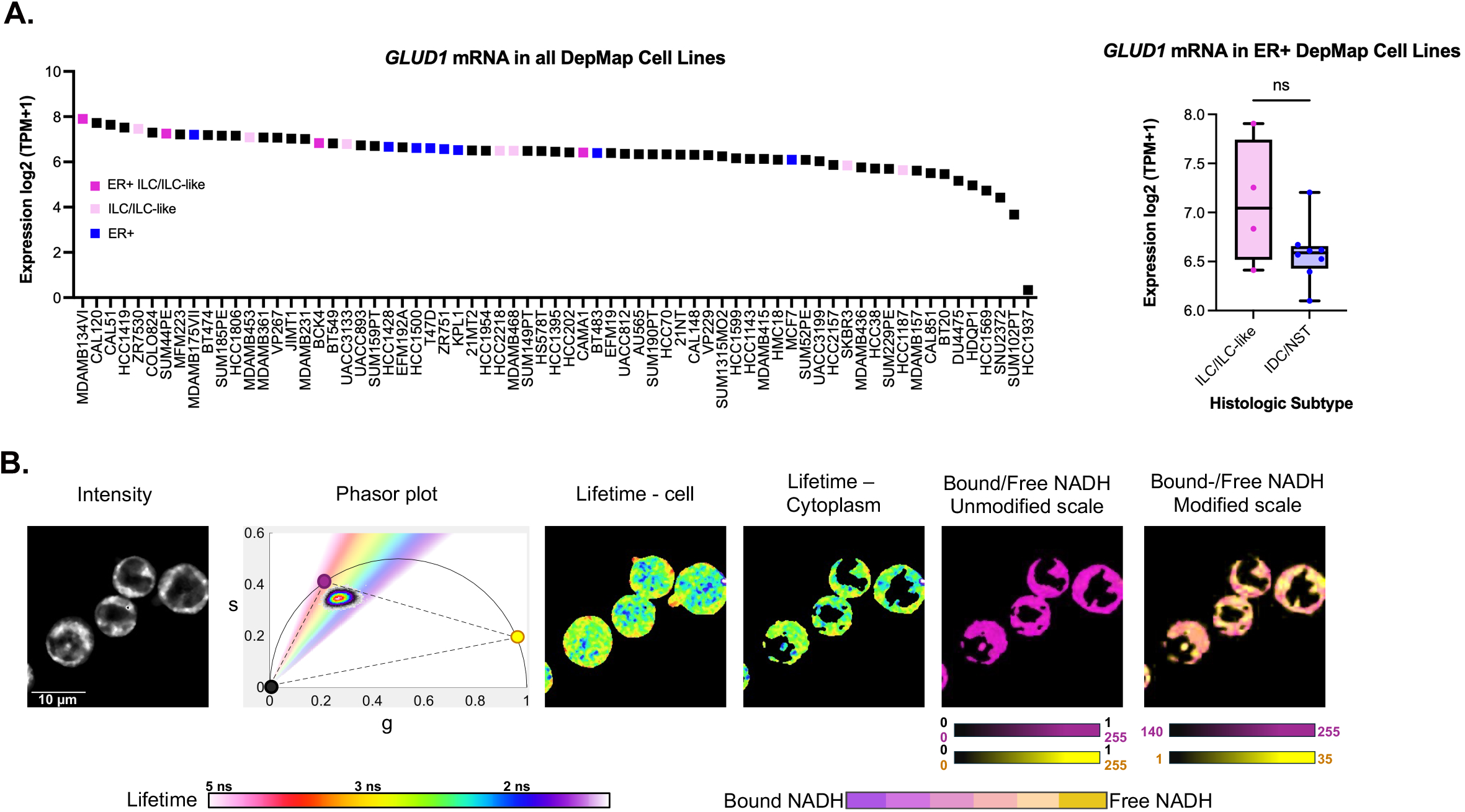
Expression *GLUD1* mRNA in ILC cell lines and phasor-FLIM metabolic imaging workflow. **A**, Waterfall plot of *GLUD1* mRNA expression data from all breast cancer cell lines in DepMap release 25Q2 (left), and boxplot of *GLUD1* mRNA expression for ER+ ILC/ILC-like vs. IDC/NST cell lines in this dataset (right, line at median, error bars minimum to maximum). Data are analyzed by unpaired t-test. Cell lines are assigned as ILC/ILC-like according to Shah et al^1^. **B**, Workflow for analyzing autofluorescence FLIM images of NADH. The fluorescence lifetime of each pixel in the intensity image (left) is transformed into a phasor plot (right). Using a phase lifetime color map (rainbow scale above the phasor plot), phasor points are color-coded and mapped back onto the original image to generate lifetime-coded autofluorescence images. These images are then restricted to the cytoplasm using intensity histogram correction. Bound/free NADH fractions are calculated using a multicomponent phasor approach, where purple indicates fully bound NADH and yellow indicates fully free NADH (binary scale: 0–255). To enhance visualization of sample differences, the color scale is adjusted to 140–255 for bound NADH (purple) and 1–35 for free NADH (yellow). These scaled images are presented in Figure 6.

---

**Supplemental Table 1:** Primary Antibody/OPAL Dye Pairings and Incubation Conditions.

|  | **Antibody 1** | **Antibody 2** | **Antibody 3** | **Antibody 4** | **Antibody 5** |
| --- | --- | --- | --- | --- | --- |
| **Antigen** | CD98 | GLUD1/2 | GPX4 | panCK | SLC7A11 |
| **Company** | Santa Cruz | Cell Signaling | Abcam | Agilent | Abcam |
| **Cat#, RRID** | Cat# sc-376815,<br>RRID:AB_2938854 | Cat# 12793,<br>RRID:AB_2750880 | Cat# ab125066,<br>RRID:AB_10973901 | Cat# M3515,<br>RRID:AB_2132885 | Cat# ab175186,<br>RRID:AB_2722749 |
| **Species** | Mouse | Rabbit | Rabbit | Mouse | Rabbit |
| **Dilution** | 1:200 | 1:200 | 1:300 | 1:300 | 1:800 |
| **Incubation Time** | 1 hr | 1 hr | 1 hr | 1 hr | 1 hr |
| **Incubation Temp.** | RT | RT | RT | RT | RT |
| **Control Tissue** | Breast Cancer | Breast Cancer | Liver Cancer | Breast Cancer | Glioma |
| **OPAL Fluor.** | 520 | 650 | 570 | 690 | 620 |
| **OPAL Conc.** | 1:150 | 1:150 | 1:400 | 1:30 | 1:300 |
| **Antigen Retrieval** | AR6 | AR6 | AR6 | AR6 | AR6 |

**Supplemental Table 2:** Literature estimates of ILC prevalence in Black women.

|  |  |  |  |
| --- | --- | --- | --- |
| Author, Year (number of Black women) | Black and Lobular (%) | Significant vs NHW? | Reference |
| ^Baquet, 2008 (nr) | nr (5.5%) | yes, ↓ | 55 |
| Elledge, 1994 (1016) | nr | no | 56 |
| Gallagher, 2022 (295) | 25 (8.5%) | yes, ↓ | 57 |
| ^Han, 2021 (7292) | 541 (7.4%) | na | 58 |
| Oesterreich, 2022 (3022) | 276 (7.6%) | yes, ↓ | 59 |
| Williams, 2018 (1393) | 126 (16%) | yes, ↓ | 60 |

n, number
NHW, non-Hispanic white
^Surveillance, Epidemiology, and End Results (SEER) data
nr, not reported
na, not applicable (Black women only)
References numbered according to main manuscript text

**Supplemental Table 3:** Comorbidity Analysis. Two-sided Fisher’s exact test was used for statistical analysis.

**A) Overall comorbidity analysis for ILC and IDC/NST**
|  | ILC | IDC | P-value |
| --- | --- | --- | --- |
| Comorbidity | 19 (26.389%) | 17 (34%) | 0.4216 (NS) |

**B) Categorical comorbidity analysis for ILC and IDC/NST**
|  | ILC | IDC | P-value |
| --- | --- | --- | --- |
| Diabetes mellitus | 5 (6.944%) | 4 (8%) | >0.9999 (NS) |
| Hypercholesterolemia and/or hyperlipidemia | 8 (11.111%) | 1 (2%) | 0.0804 (NS) |
| Obesity | 0 (0%) | 7 (14%) | 0.0015 (**p<0.01) |
| Hypertension | 19 (26.389%) | 12 (24%) | 0.8345 (NS) |
| Chronic kidney disease | 0 (0%) | 1 (2%) | 0.4050 (NS) |

## References

1. Christgen, M. et al. Lobular Breast Cancer: Histomorphology and Different Concepts of a Special Spectrum of Tumors. Cancers 13, 3695 (2021).

2. McCart Reed, A. E., Kalinowski, L., Simpson, P. T. & Lakhani, S. R. Invasive lobular carcinoma of the breast: the increasing importance of this special subtype. Breast Cancer Res 23, 6 (2021).

3. Ciriello, G. et al. Comprehensive Molecular Portraits of Invasive Lobular Breast Cancer. Cell 163, 506–19 (2015).

4. Ulaner, G. A. & Schuster, D. M. Amino Acid Metabolism as a Target for Breast Cancer Imaging. PET Clin 13, 437–444 (2018).

5. Savir-Baruch, B., Zanoni, L. & Schuster, D. M. Imaging of Prostate Cancer Using Fluciclovine. PET Clinics 12, 145–157 (2017).

6. Ulaner, G. A. et al. Initial Results of a Prospective Clinical Trial of 18F-Fluciclovine PET/CT in Newly Diagnosed Invasive Ductal and Invasive Lobular Breast Cancers. J. Nucl. Med. 57, 1350–1356 (2016).

7. Ulaner, G. A. et al. Prospective Clinical Trial of 18F-Fluciclovine PET/CT for Determining the Response to Neoadjuvant Therapy in Invasive Ductal and Invasive Lobular Breast Cancers. J. Nucl. Med. 58, 1037–1042 (2017).

8. Mushtaq, A. et al. Prospective investigation of amino acid transport and PSMA-targeted positron emission tomography for metastatic lobular breast carcinoma. Eur J Nucl Med Mol Imaging 10.1007/s00259-024-06830-7 (2024) doi:10.1007/s00259-024-06830-7.

9. Du, T. et al. Key regulators of lipid metabolism drive endocrine resistance in invasive lobular breast cancer. Breast Cancer Res. 20, 106 (2018).

10. Du, T. et al. Invasive lobular and ductal breast carcinoma differ in immune response, protein translation efficiency and metabolism. Sci Rep 8, 7205 (2018).

11. Cha, Y. J., Kim, H. M. & Koo, J. S. Expression of Lipid Metabolism-Related Proteins Differs between Invasive Lobular Carcinoma and Invasive Ductal Carcinoma. International Journal of Molecular Sciences 18, 232 (2017).

12. Sottnik, J. L. et al. WNT4 Regulates Cellular Metabolism via Intracellular Activity at the Mitochondria in Breast and Gynecologic Cancers. Cancer Res Commun 4, 134–151 (2024).

13. Stires, H. et al. Integrated molecular analysis of Tamoxifen-resistant invasive lobular breast cancer cells identifies MAPK and GRM/mGluR signaling as therapeutic vulnerabilities. Mol Cell Endocrinol 471, 105–117 (2018).

14. Olukoya, A. O., et al. Riluzole Suppresses Growth and Enhances Response to Endocrine Therapy in ER+ Breast Cancer. J Endocr Soc 7, bvad117 (2023).

15. Bahnassy, S., Sikora, M. J. & Riggins, R. B. Unlocking the Mysteries of Lobular Breast Cancer Biology Needs the Right Combination of Preclinical Models. Mol Cancer Res 20, 837–840 (2022).

16. Sikora, M. J. & Ostrander, J. H. A Path to Precision Metabolic Treatment in Breast Cancer: Riluzole, Glutamate Signaling, and Invasive Lobular Carcinoma. J Endocr Soc 8, bvad171 (2024).

17. Ma, Z. et al. Glutamate dehydrogenase 1: A novel metabolic target in inhibiting acute myeloid leukaemia progression. Br J Haematol 202, 566–577 (2023).

18. Spinelli, J. B. et al. Metabolic recycling of ammonia via glutamate dehydrogenase supports breast cancer biomass. Science 358, 941–946 (2017).

19. Liu, J., Xia, X. & Huang, P. xCT: A Critical Molecule That Links Cancer Metabolism to Redox Signaling. Molecular Therapy 28, 2358–2366 (2020).

20. Shin, S.-S. et al. Participation of xCT in melanoma cell proliferation in vitro and tumorigenesis in vivo. Oncogenesis 7, 86 (2018).

21. Xia, P. & Dubrovska, A. CD98 heavy chain as a prognostic biomarker and target for cancer treatment. Front. Oncol. 13, (2023).

22. El Ansari, R., et al. The multifunctional solute carrier 3A2 (SLC3A2) confers a poor prognosis in the highly proliferative breast cancer subtypes. Br J Cancer 118, 1115–1122 (2018).

23. Ye, L. et al. Metabolism-regulated ferroptosis in cancer progression and therapy. Cell Death Dis 15, 1–12 (2024).

24. Okazaki, S. et al. Glutaminolysis-related genes determine sensitivity to xCT-targeted therapy in head and neck squamous cell carcinoma. Cancer Sci 110, 3453–3463 (2019).

25. Zhang, X. et al. Endogenous glutamate determines ferroptosis sensitivity via ADCY10-dependent YAP suppression in lung adenocarcinoma. Theranostics 11, 5650–5674 (2021).

26. Liu, J. et al. An Integrated TCGA Pan-Cancer Clinical Data Resource to Drive High-Quality Survival Outcome Analytics. Cell 173, 400–416.e11 (2018).

27. Pereira, B. et al. The somatic mutation profiles of 2,433 breast cancers refines their genomic and transcriptomic landscapes. Nat Commun 7, 11479 (2016).

28. Saal, L. H. et al. The Sweden Cancerome Analysis Network - Breast (SCAN-B) Initiative: a large-scale multicenter infrastructure towards implementation of breast cancer genomic analyses in the clinical routine. Genome Med 7, 20 (2015).

29. Brueffer, C. et al. The mutational landscape of the SCAN-B real-world primary breast cancer transcriptome. EMBO Mol Med 12, e12118 (2020).

30. Cerami, E. et al. The cBio cancer genomics portal: an open platform for exploring multidimensional cancer genomics data. Cancer Discov 2, 401–4 (2012).

31. Gao, J. et al. Integrative analysis of complex cancer genomics and clinical profiles using the cBioPortal. Sci Signal 6, pl1 (2013).

32. Sottnik, J. L. et al. Mediator of DNA Damage Checkpoint 1 (MDC1) Is a Novel Estrogen Receptor Coregulator in Invasive Lobular Carcinoma of the Breast. Mol Cancer Res 19, 1270–1282 (2021).

33. Fernandez, A. I. et al. The orphan nuclear receptor estrogen-related receptor beta (ERRβ) in triple-negative breast cancer. Breast Cancer Research and Treatment 179, 585–604 (2020).

34. Riggins, R. B. et al. ERRgamma Mediates Tamoxifen Resistance in Novel Models of Invasive Lobular Breast Cancer. Cancer Res. 68, 8908–8917 (2008).

35. Sikora, M. J. et al. WNT4 mediates estrogen receptor signaling and endocrine resistance in invasive lobular carcinoma cell lines. Breast Cancer Res 18, 92 (2016).

36. Schindelin, J., et al. Fiji: an open-source platform for biological-image analysis. Nature Methods 9, 676–682 (2012).

37. Vallmitjana, A. et al. GSLab: open-source platform for advanced phasor analysis in fluorescence microscopy. Bioinformatics 41, btaf162 (2025).

38. Malacrida, L., Ranjit, S., Jameson, D. M. & Gratton, E. The Phasor Plot: A Universal Circle to Advance Fluorescence Lifetime Analysis and Interpretation. Annu Rev Biophys 50, 575–593 (2021).

39. Ranjit, S., Malacrida, L., Jameson, D. M. & Gratton, E. Fit-free analysis of fluorescence lifetime imaging data using the phasor approach. Nat Protoc 13, 1979–2004 (2018).

40. Stringari, C. et al. Metabolic trajectory of cellular differentiation in small intestine by Phasor Fluorescence Lifetime Microscopy of NADH. Sci Rep 2, 568 (2012).

41. Vallmitjana, A. et al. Resolution of 4 components in the same pixel in FLIM images using the phasor approach. Methods Appl. Fluoresc. 8, 035001 (2020).

42. Reinhart, G. D., Marzola, P., Jameson, D. M. & Gratton, E. A method for on-line background subtraction in frequency domain fluorometry. J Fluoresc 1, 153–162 (1991).

43. Ranjit, S., Malacrida, L., Stakic, M. & Gratton, E. Determination of the metabolic index using the fluorescence lifetime of free and bound nicotinamide adenine dinucleotide using the phasor approach. J Biophotonics 12, e201900156 (2019).

44. Giaquinto, A. N., Freedman, R. A., Newman, L. A., Jemal, A. & Siegel, R. L. Lobular breast cancer statistics, 2025. Cancer 131, e70061 (2025).

45. Quinn, R. M., Bernal, A. M., Oh, S. Y. & Anampa, J. D. Trends in Incidence of Invasive Lobular Carcinoma of the Breast by Race: Patterns by Age, Cancer Stage, and Socioeconomic Factors in the United States, 1992-2019. Clinical Breast Cancer 10.1016/j.clbc.2024.12.015 (2024) doi:10.1016/j.clbc.2024.12.015.

46. Sandhu, S., Lim, D. W. & Giannakeas, V. Racial disparities in presentation and survival for lobular breast cancer. J Clin Oncol 42, 1596–1596 (2024).

47. Sreekumar, S. et al. Differential regulation and targeting of estrogen receptor α turnover in invasive lobular breast carcinoma. Endocrinology 10.1210/endocr/bqaa109 (2020) doi:10.1210/endocr/bqaa109.

48. Timmerman, L. A. et al. Glutamine sensitivity analysis identifies the xCT antiporter as a common triple-negative breast tumor therapeutic target. Cancer Cell 24, 450–465 (2013).

49. Coloff, J. L. et al. Differential Glutamate Metabolism in Proliferating and Quiescent Mammary Epithelial Cells. Cell Metab 23, 867–880 (2016).

50. Michaelson, J. S. et al. Predicting the survival of patients with breast carcinoma using tumor size. Cancer 95, 713–723 (2002).

51. Parab, A. Z. et al. Socioecologic Factors and Racial Differences in Breast Cancer Multigene Prognostic Scores in US Women. JAMA Network Open 7, e244862 (2024).

52. Van Alsten, S. C. et al. Differences in 21-Gene and PAM50 Recurrence Scores in Younger and Black Women With Breast Cancer. JCO Precis Oncol e2400137 (2024) doi:10.1200/PO.24.00137.

53. Rauscher, G. H. et al. Racial disparity in survival from estrogen and progesterone receptor positive breast cancer: implications for reducing breast cancer mortality disparities. Breast cancer research and treatment 163, 321 (2017).

54. Torres, J. M., Sodipo, M. O., Hopkins, M. F., Chandler, P. D. & Warner, E. T. Racial Differences in Breast Cancer Survival Between Black and White Women According to Tumor Subtype: A Systematic Review and Meta-Analysis. J Clin Oncol JCO2302311 (2024) doi:10.1200/JCO.23.02311.

55. Baquet, C. R., Mishra, S. I., Commiskey, P., Ellison, G. L. & DeShields, M. Breast cancer epidemiology in blacks and whites: disparities in incidence, mortality, survival rates and histology. J Natl Med Assoc 100, 480–488 (2008).

56. Elledge, R. M., Clark, G. M., Chamness, G. C. & Osborne, C. K. Tumor Biologic Factors and Breast Cancer Prognosis Among White, Hispanic, and Black Women in the United States 1. JNCI: Journal of the National Cancer Institute 86, 705–712 (1994).

57. Gallagher, E. J. et al. Insulin resistance and racial disparities in breast cancer prognosis: a multi-center cohort study. 10.1530/ERC-22-0106 (2022) doi:10.1530/ERC-22-0106.

58. Han, Y. et al. Breast Cancer Mortality Hot Spots Among Black Women With de Novo Metastatic Breast Cancer. JNCI Cancer Spectrum 5, pkaa086 (2021).

59. Oesterreich, S. et al. Clinicopathological Features and Outcomes Comparing Patients With Invasive Ductal and Lobular Breast Cancer. JNCI: Journal of the National Cancer Institute 114, 1511–1522 (2022).

60. Williams, L. A. et al. Reproductive risk factor associations with lobular and ductal carcinoma in the Carolina Breast Cancer Study. Cancer Causes Control 29, 25–32 (2018).

61. De Schepper, M. et al. Integration of Pathological Criteria and Immunohistochemical Evaluation for Invasive Lobular Carcinoma Diagnosis: Recommendations From the European Lobular Breast Cancer Consortium. Mod Pathol 37, 100497 (2024).

62. Djerroudi, L. et al. E-Cadherin Mutational Landscape and Outcomes in Breast Invasive Lobular Carcinoma. Mod Pathol 37, 100570 (2024).

63. Xie, Z. et al. Gene Set Knowledge Discovery with Enrichr. Current Protocols 1, e90 (2021).

64. Kuleshov, M. V. et al. Enrichr: a comprehensive gene set enrichment analysis web server 2016 update. Nucleic Acids Res 44, W90–97 (2016).

65. Chen, E. Y. et al. Enrichr: interactive and collaborative HTML5 gene list enrichment analysis tool. BMC Bioinformatics 14, 128 (2013).

66. Oikari, S. et al. UDP-sugar accumulation drives hyaluronan synthesis in breast cancer. Matrix Biol 67, 63–74 (2018).

67. Niu, F. et al. Arginase: An emerging and promising therapeutic target for cancer treatment. Biomedicine & Pharmacotherapy 149, 112840 (2022).

68. Nyrop, K. A. et al. Obesity, comorbidities, and treatment selection in Black and White women with early breast cancer. Cancer 127, 922–930 (2021).

69. Han, H. et al. Hypertension and breast cancer risk: a systematic review and meta-analysis. Sci Rep 7, 44877 (2017).

70. Anwar, S. L. et al. Metabolic comorbidities and the association with risks of recurrent metastatic disease in breast cancer survivors. BMC Cancer 21, 590 (2021).

71. Monzavi-Karbassi, B. et al. Pre-diagnosis blood glucose and prognosis in women with breast cancer. Cancer & Metabolism 4, 7 (2016).

72. Arafeh, R., Shibue, T., Dempster, J. M., Hahn, W. C. & Vazquez, F. The present and future of the Cancer Dependency Map. Nat Rev Cancer 25, 59–73 (2025).

73. Shah, O. S. et al. Multi-omic characterization of ILC and ILC-like cell lines as part of ILC cell line encyclopedia (ICLE) defines new models to study potential biomarkers and explore therapeutic opportunities. 2023.09.26.559548 Preprint at 10.1101/2023.09.26.559548 (2023).

74. Su, X. B. & Pillus, L. Functions for diverse metabolic activities in heterochromatin. Proceedings of the National Academy of Sciences 113, E1526–E1535 (2016).

75. Traube, F. R. et al. Redirected nuclear glutamate dehydrogenase supplies Tet3 with α-ketoglutarate in neurons. Nat Commun 12, 4100 (2021).

76. Mohammed, H. et al. Endogenous purification reveals GREB1 as a key estrogen receptor regulatory factor. Cell Rep 3, 342–349 (2013).

77. Mi, W. et al. BET inhibition induces GDH1-dependent glutamine metabolic remodeling and vulnerability in liver cancer. Life Metabolism 3, loae016 (2024).

78. Kang, J. et al. EGFR-phosphorylated GDH1 harmonizes with RSK2 to drive CREB activation and tumor metastasis in EGFR-activated lung cancer. Cell Rep 41, 111827 (2022).

79. Jin, L. et al. Glutamate dehydrogenase 1 signals through antioxidant glutathione peroxidase 1 to regulate redox homeostasis and tumor growth. Cancer Cell 27, 257–270 (2015).

80. Wang, Q. et al. Therapeutic targeting of glutamate dehydrogenase 1 that links metabolic reprogramming and Snail-mediated epithelial-mesenchymal transition in drug-resistant lung cancer. Pharmacol Res 185, 106490 (2022).

81. Ranjit, S., Datta, R., Dvornikov, A. & Gratton, E. Multicomponent Analysis of Phasor Plot in a Single Pixel to Calculate Changes of Metabolic Trajectory in Biological Systems. J Phys Chem A 123, 9865–9873 (2019).

82. Vidyadharan, V. A., Betancourt, A., Smith, C., Blesson, C. S. & Yallampalli, C. Maternal Low-Protein Diet Leads to Mitochondrial Dysfunction and Impaired Energy Metabolism in the Skeletal Muscle of Male Rats. International Journal of Molecular Sciences 25, 12860 (2024).

83. dos Santos, M. P., et al. Lipopolysaccharide Induces Mitochondrial Fragmentation and Energetic Shift in Reactive Microglia: Evidence for a Cell-Autonomous Program of Metabolic Plasticity. Toxins 17, 293 (2025).

84. Hershberger, K. A. et al. Early-life mitochondrial DNA damage results in lifelong deficits in energy production mediated by redox signaling in *Caenorhabditis elegans*. Redox Biology 43, 102000 (2021).

85. Cao, J. et al. Targeting estrogen-regulated system xc- promotes ferroptosis and endocrine sensitivity of ER+ breast cancer. Cell Death Dis 16, 30 (2025).

86. Crocco, P., Dato, S., La Grotta, R., Passarino, G. & Rose, G. Evidence for a relationship between genetic polymorphisms of the L-DOPA transporter LAT2/4F2hc and risk of hypertension in the context of chronic kidney disease. BMC Med Genomics 17, 163 (2024).

87. Bajaj, J. et al. CD98-Mediated Adhesive Signaling Enables the Establishment and Propagation of Acute Myelogenous Leukemia. Cancer Cell 30, 792–805 (2016).

88. Kawasaki, Y., Omori, Y., Suzuki, S. & Yamada, T. CD98hc as a marker of radiotherapy-resistant cancer stem cells in head and neck squamous cell carcinoma. Arch Med Sci 19, 1859–1868 (2023).

89. Saito, Y. et al. Polarity protein SCRIB interacts with SLC3A2 to regulate proliferation and tamoxifen resistance in ER+ breast cancer. Commun Biol 5, 1–10 (2022).

90. Cormerais, Y. et al. Genetic Disruption of the Multifunctional CD98/LAT1 Complex Demonstrates the Key Role of Essential Amino Acid Transport in the Control of mTORC1 and Tumor Growth. Cancer Research 76, 4481–4492 (2016).

91. Boulter, E. et al. Cell metabolism regulates integrin mechanosensing via an SLC3A2-dependent sphingolipid biosynthesis pathway. Nat Commun 9, 4862 (2018).

92. Wu, J. et al. Intercellular interaction dictates cancer cell ferroptosis via NF2-YAP signalling. Nature 572, 402–406 (2019).

93. Panzilius, E. et al. Cell density-dependent ferroptosis in breast cancer is induced by accumulation of polyunsaturated fatty acid-enriched triacylglycerides. bioRxiv 417949 (2019) doi:10.1101/417949.

94. Hu, K. et al. Efficacy of FERscore in predicting sensitivity to ferroptosis inducers in breast cancer. NPJ Breast Cancer 10, 74 (2024).

95. Xu, Z. et al. RelB-activated GPX4 inhibits ferroptosis and confers tamoxifen resistance in breast cancer. Redox Biol 68, 102952 (2023).

96. Minikes, A. M. et al. E-cadherin is a biomarker for ferroptosis sensitivity in diffuse gastric cancer. Oncogene 42, 848–857 (2023).

97. Mbah, N. E. & Lyssiotis, C. A. Metabolic regulation of ferroptosis in the tumor microenvironment. Journal of Biological Chemistry 298, (2022).

98. Liu, Y., Duan, C., Dai, R. & Zeng, Y. Ferroptosis-mediated Crosstalk in the Tumor Microenvironment Implicated in Cancer Progression and Therapy. Front. Cell Dev. Biol. 9, (2021).

99. Craze, M. L. et al. Glutamate dehydrogenase (GLUD1) expression in breast cancer. Breast Cancer Res Treat 174, 79–91 (2019).

100. Shah, O. S. et al. Spatial molecular profiling of mixed invasive ductal and lobular breast cancers reveals heterogeneity in intrinsic molecular subtypes, oncogenic signatures, and mutations. Proceedings of the National Academy of Sciences 121, e2322068121 (2024).

101. Shackleford, M. T. et al. Estrogen Regulation of mTOR Signaling and Mitochondrial Function in Invasive Lobular Carcinoma Cell Lines Requires WNT4. Cancers 12, 2931 (2020).

102. Sottnik, J. L., et al. Co-regulator activity of Mediator of DNA Damage Checkpoint 1 (MDC1) is associated with DNA repair dysfunction and PARP inhibitor sensitivity in lobular carcinoma of the breast. bioRxiv 2023.10.29.564555 (2025) doi:10.1101/2023.10.29.564555.

103. Hensel, A., Váraljai, R. & Knauer, S. K. Raising the iron curtain: Lactate’s secret role in oxidative stress defense. Redox Biology 85, 103754 (2025).

104. Santaliz-Casiano, A. et al. Identification of metabolic pathways contributing to ER+ breast cancer disparities using a machine-learning pipeline. Sci Rep 13, 12136 (2023).

105. Bobba-Alves, N. et al. Cellular allostatic load is linked to increased energy expenditure and accelerated biological aging. Psychoneuroendocrinology 155, 106322 (2023).

106. Lee, M. et al. Assessing the Correlation between Allostatic Load and False-Positive Image-Guided Breast Biopsies. J Womens Health (Larchmt) 10.1089/jwh.2024.0039 (2024) doi:10.1089/jwh.2024.0039.

107. Wang, F. et al. Allostatic load and risk of invasive breast cancer among postmenopausal women in the U.S. Preventive Medicine 178, 107817 (2024).

108. Kaur, M. et al. Area Deprivation Index in Patients with Invasive Lobular Carcinoma of the Breast: Associations with Tumor Characteristics and Outcomes. Cancer Epidemiology, Biomarkers & Prevention 32, 1107–1113 (2023).

## Supplement References

1. Shah, O. S. et al. Multi-omic characterization of ILC and ILC-like cell lines as part of ILC cell line encyclopedia (ICLE) defines new models to study potential biomarkers and explore therapeutic opportunities. 2023.09.26.559548 Preprint at 10.1101/2023.09.26.559548 (2023).

